# Layer-specific, retinotopically-diffuse modulation in human visual cortex by emotional faces

**DOI:** 10.1101/2022.07.10.499443

**Authors:** Tina T. Liu, Jason Z Fu, Yuhui Chai, Shruti Japee, Gang Chen, Leslie G. Ungerleider, Elisha P. Merriam

**Affiliations:** Sections on Neurocircuitry, National Institute of Mental Health, NIH, Bethesda, MD, USA; Functional Imaging Methods, National Institute of Mental Health, NIH, Bethesda, MD, USA; Laboratory of Brain and Cognition, National Institute of Mental Health, NIH, Bethesda, MD, USA; Scientific and Statistical Computing Core, National Institute of Mental Health, NIH, Bethesda, MD, USA

**Keywords:** layer-specific fMRI, emotion, primary visual cortex, amygdala

## Abstract

Emotionally expressive faces evoke enhanced neural responses in multiple brain regions, a phenomenon thought to depend critically on the amygdala. This emotion-related modulation is evident even in primary visual cortex (V1), providing a potential neural substrate by which emotionally salient stimuli can affect perception. How does emotional valence information, computed in the amygdala, reach V1? Here we use high-resolution functional MRI to investigate the layer profile and retinotopic distribution of neural activity specific to emotional facial expressions. Across three experiments, human participants viewed centrally presented face stimuli varying in emotional expression and performed a gender judgment task. We found that facial valence sensitivity was evident only in superficial cortical layers and was not restricted to the retinotopic location of the stimuli, consistent with diffuse feedback-like projections from the amygdala. Together, our results provide a feedback mechanism by which the amygdala directly modulates activity at the earliest stage of visual processing.

## Main

Emotional facial expressions convey a wealth of non-verbal information, such as an individual’s mood, state of mind and intention, and hence are critically important for social communication. In macaques, emotionally expressive faces compared to neutral faces elicit greater responses in the amygdala and face-selective patches^1,2^, a phenomenon commonly referred to as the “valence effect”^1^. In humans, fearful faces evoke stronger blood oxygen level dependent (BOLD) activity than neutral faces in the amygdala^3^, face-selective cortex^4^, and V1^5^. Despite evidence from neuroanatomy^6,7^, neuroimaging^5,8^, and neuropsychology^9,10^ suggesting the amygdala plays a role in coordinating how we respond to biologically relevant stimuli^11^, especially fearful faces^12^, the presence of valence effect in V1 is surprising because early visual cortex is not typically thought to be sensitive to emotional aspects of visual stimuli. The valence effect in the visual cortex is diminished in human patients^13^ and monkeys^14^ with amygdala lesions, suggesting that feedback from the amygdala is the source of the valence effect in V1. However, functional pathways by which emotional information is transmitted from the amygdala to V1 remains unclear.

Anatomical studies in non-human primates have demonstrated an asymmetric pattern of connectivity between the amygdala and visual cortex^7,15^. That is, the lateral nucleus of the amygdala receives feedforward inputs propagated from V1 to IT cortex, while the basal nucleus of the amygdala sends widespread projections to areas all along the ventral visual pathway. Thus, one possibility is that valence information computed in the amygdala reaches V1 via intracortical feedback projections from higher-order visual regions such as the fusiform face area (FFA). A second possibility is that valence information reaches V1 via direct anatomical projections from the basal amygdala^15,16^. These two competing hypotheses make different predictions regarding both the laminar profile and the retinotopic specificity of activity in V1. Feedback projections from higher-order visual areas terminate in superficial and deep layers of V1^17^. Moreover, feedback projections from higher-order visual areas are thought to either be retinotopically specific, or favor the foveal representation^18–20^. In contrast, feedback projections from the amygdala terminate exclusively in the superficial layers of V1 in macaque monkeys, and are not topographic, present throughout the entire extent of V1^15^. We used a facial expression protocol that has been widely used to study emotional responses in both humans^21^ and monkeys^1^ and evaluated which of these two hypotheses most closely matched the pattern of neural activity that we measured with functional magnetic resonance imaging (fMRI). We conducted three experiments at two field strengths (7T and 3T) while human participants (*n*=25, number of scan sessions=43, see Table 1) viewed face stimuli blocked by emotional expressions, and performed a gender judgment task orthogonal to emotional expression.

**Table 1.** Demographics and scan details of the healthy volunteers (number of unique participants = 25, total scan sessions = 43).

| Participant | Gender | Age | Number of scans acquired | 3T BOLD | 7T BOLD | 7T VASO | Face localizer acquired |
| --- | --- | --- | --- | --- | --- | --- | --- |
| 1 | F | 23 | 3 | 1 | 1 | 1 | Y |
| 2 | F | 23 | 1 | 0 | 1 | 0 | Y |
| 3 | F | 29 | 2 | 1 | 1 | 0 | Y |
| 4 | F | 27 | 4 | 1 | 1 | 2 | Y |
| 5 | F | 23 | 1 | 1 | 0 | 0 | N |
| 6 | F | 23 | 1 | 1 | 0 | 0 | N |
| 7 | F | 23 | 1 | 1 | 0 | 0 | N |
| 8 | M | 24 | 1 | 0 | 1 | 0 | N |
| 9 | M | 23 | 2 | 1 | 1 | 0 | Y |
| 10 | M | 31 | 1 | 0 | 0 | 1 | N |
| 11 | M | 23 | 1 | 1 | 0 | 0 | Y |
| 12 | M | 24 | 1 | 0 | 1 | 0 | Y |
| 13 | F | 22 | 1 | 1 | 0 | 0 | Y |
| 14 | M | 31 | 3 | 1 | 0 | 2 | Y |
| 15 | M | 22 | 4 | 1 | 1 | 2 | Y |
| 16 | F | 21 | 1 | 0 | 1 | 0 | Y |
| 17 | M | 34 | 2 | 1 | 1 | 0 | Y |
| 18 | M | 42 | 2 | 1 | 1 | 0 | Y |
| 19 | M | 23 | 1 | 0 | 1 | 0 | N |
| 20 | M | 25 | 3 | 1 | 1 | 1 | Y |
| 21 | F | 24 | 1 | 0 | 0 | 1 | N |
| 22 | F | 25 | 2 | 0 | 0 | 2 | N |
| 23 | M | 37 | 2 | 0 | 0 | 2 | N |
| 24 | F | 23 | 1 | 0 | 0 | 1 | N |
| 25 | M | 22 | 1 | 0 | 1 | 0 | Y |
| Total number of scans per experiment: |  |  |  | 14 | 14 | 15 |  |

We found a robust valence effect in the BOLD fMRI measurements, present in many brain regions, including V1, replicating previous reports^5,22^. We then performed an inter-area correlation analysis revealing that the amygdala is the source of the widespread valence effect in visual cortex. To further understand the mechanisms by which the amygdala modulates responses in V1, we used vascular-space-occupancy (VASO) fMRI at 7T to measure changes in cerebral blood volume (CBV) across cortical layers^23,24^. We found that the valence effect in V1 was only evident in superficial cortical depths. Retinotopic analysis revealed that the valence effect was present throughout all of V1, including portions of V1 that were not stimulated by the face stimuli. Together, our results demonstrate a mechanism of facial valence modulation—valence information computed in the amygdala is fed back to V1 via direct anatomical projections to enhance the processing of low-level stimulus features associated with fear-inducing stimuli.

## Results

### Widely distributed valence effect

Face stimuli with fearful or happy expressions evoked a larger BOLD response than faces with neutral expressions in nearly every retinotopically-defined cortical area (Fig. 1b-c), a phenomenon referred to as the valence effect^1^. The valence effect was highly reliable. We observed the valence effect at both 3T and 7T field strengths, in both the group results (Fig. 1b-c) and in individual participants (Supplementary Figs. 1-3). To quantify the valence effect, we segmented the visual cortex into 13 regions of interest (ROIs, labeled in Fig. 1b-c) using a probabilistic retinotopic atlas^25^ and functionally defined the amygdala and the FFA using an independent localizer experiment (see Methods). We then averaged BOLD activity across visually-responsive voxels within each area, also averaging responses across sessions for those participants who were scanned in multiple sessions (see Table 1). We used a Bayesian multilevel (BML) modeling approach to derive a robust estimate of the strength of the valence effects, which we plot as a negative valence index and as a positive valence index in each ROI (Fig. 1d-e). The negative valence effect was evident in every visual area, including the amygdala and V1 (Fig. 1b&d). We also observed reliable, albeit less pronounced, positive valence effects associated with happy facial expressions in many visual areas (Fig. 1c&e).

**Fig. 1:**
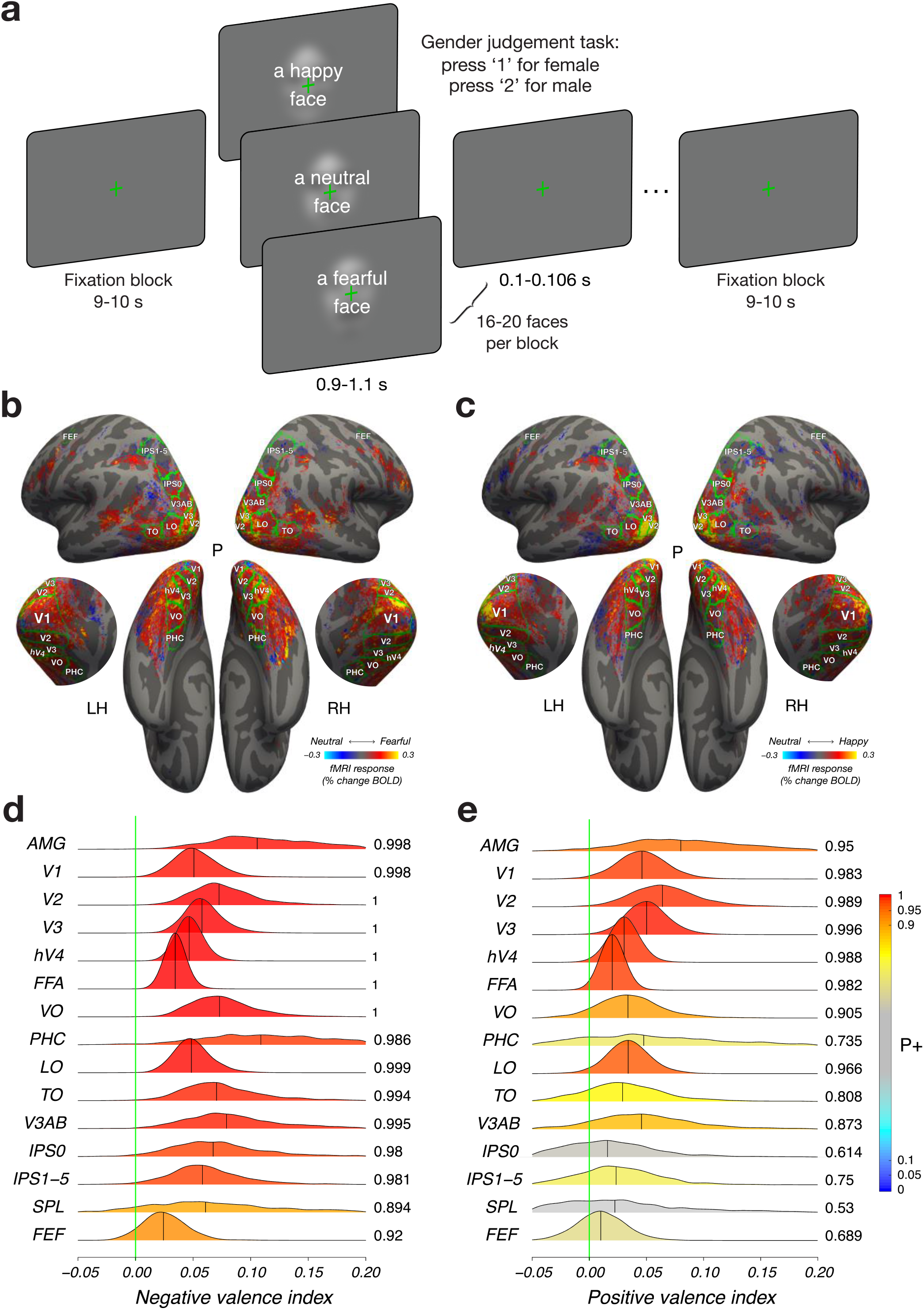
Distribution of valence modulation throughout the brain. **a**, Stimuli and experimental protocol. Participants viewed a series of closely cropped face stimuli balanced for low-level visual properties and blocked by emotional expression (happy, neutral, fearful) while performing a gender judgment task orthogonal to emotional expression. **b**, Fearful faces evoked a larger response than neutral faces in nearly every area that exhibited visual responses. Hue indicates subtraction of response amplitude to neutral from fearful faces for each voxel that exhibited a reliable visual response (coefficient of determination R^2^>0.1) in at least one third of the participants. **c**, Happy faces evoked a numerically larger response than neutral faces in many areas that exhibited visual responses. Hue indicates subtraction of response amplitude to happy faces and neutral faces for each voxel that exhibited a reliable visual response (coefficient of determination R^2^>0.1) in at least one third of the participants. **b-c**, Lateral (top), medial (middle), and ventral (bottom) views of the freesurfer average cortical surface template^26^. Green lines, areal boundaries from probabilistic retinotopic atlas^25^. **d**, Posterior distribution of negative valence effect (fearful versus neutral index: 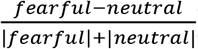) in each ROI. **e**, Posterior distribution of the positive valence effect (happy versus neutral index: 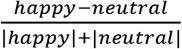) in each ROI. **d-e**, Hue indicates the strength of statistical evidence according to the Bayesian Multilevel (BML) model^27^ (see Methods), shown through *P*+ (value at right side of each posterior distribution), the posterior probability of each region’s effect being positive conditioning on the adopted BML model and the current data. The vertical green line indicates zero effect. ROIs with strong evidence of the valence effect can be identified as the extent of the green line being farther into the tail of the posterior distribution. **b-e**, Number of unique participants scanned at 3T BOLD and 7T BOLD who were also scanned in the face localizer experiment: *n*=15 (see Table 1).

### Correlation between amygdala and visual cortex enhanced by fearful faces

We performed an inter-area correlation analysis to test whether the widespread valence effect (Fig. 1b-e) is due to input from the amygdala, or, alternatively, to pervasive cortico-cortical interactions. We characterized changes in intrinsic activity fluctuations that were not directly induced by the stimulus^28^ by removing (i.e., regressing out) the stimulus-driven component of the fMRI BOLD time series (Fig. 2a, orange line) from the measured response time series (Fig. 2a, green line) averaged across voxels within each ROI. This procedure produced a residual time series (Fig. 2a, purple line), separately for each ROI and for each participant, that were then used to construct correlation matrices between each pair of ROIs. Three matrices were constructed: one matrix corresponding to epochs of fearful faces, one corresponding to epochs of happy faces, and one corresponding to epochs of neutral faces. Finally, we computed the negative valence effect by subtracting the neutral correlation matrix (Fig. 2b, middle) from the fearful correlation matrix (Fig. 2b, left), and the positive valence effect by subtracting the neutral correlation matrix (Fig. 2c, middle) from the happy correlation matrix (Fig. 2c, left). If valence information reaches V1 via intracortical feedback projections, intrinsic fluctuations between V1 and adjacent extrastriate areas, such as V2 or V3, should be higher in the fearful than in the neutral condition. In contrast, if valence information reaches V1 via direct anatomical projections from the basal amygdala, the intrinsic fluctuations between V1 and the amygdala should be higher in the fearful than in the neutral face condition.

**Fig. 2:**
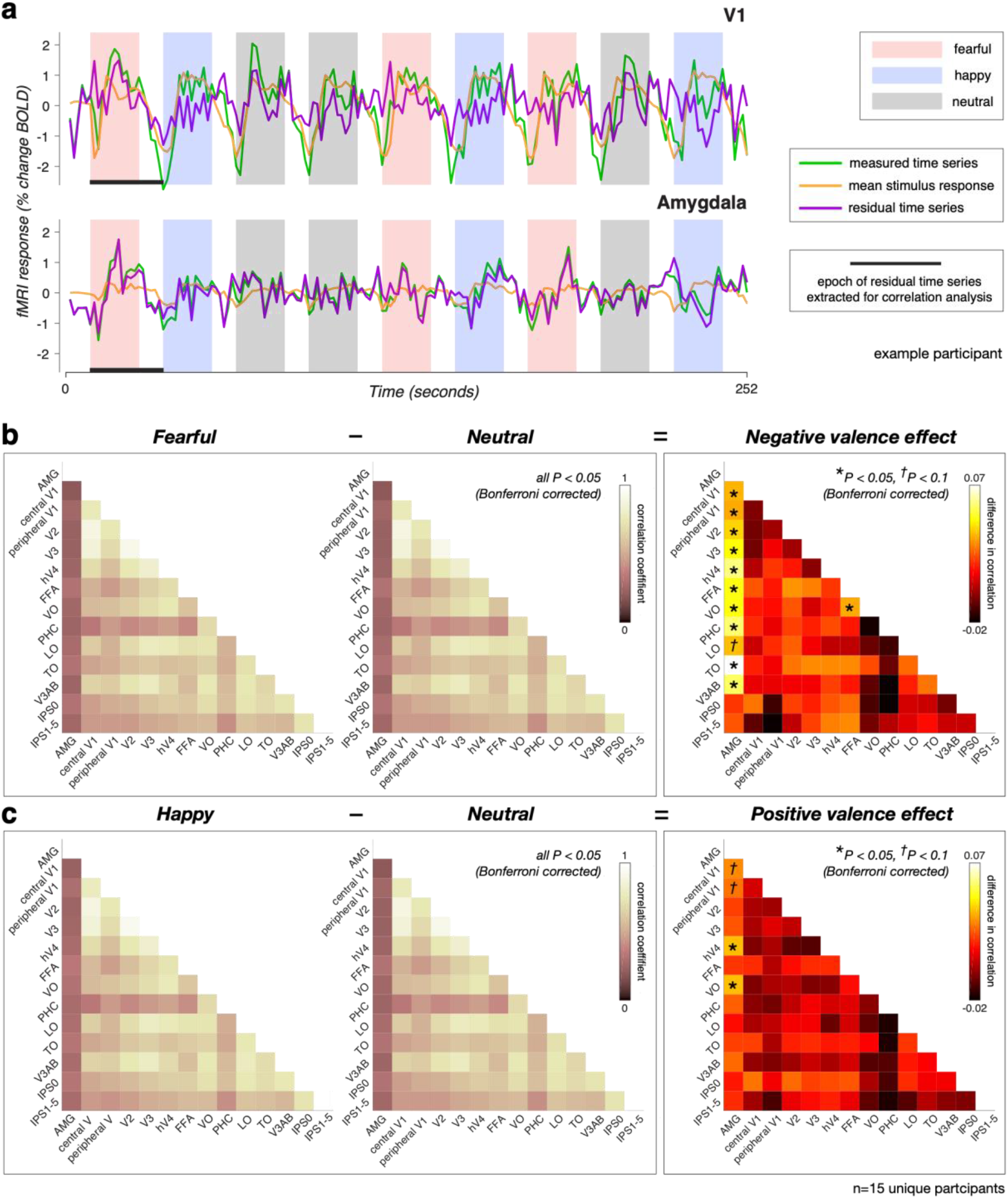
Inter-area correlation reveals enhanced interactions with amygdala when viewing fearful faces. **a**, fMRI time series from V1 (top) and the amygdala (bottom) from a single run from an example participant in the 7T BOLD experiment, consisting of three 18 s blocks of trials of each facial expression (pink: fearful; blue: happy; gray: neutral) with interleaved blocks of fixation of 9 s. Three time series are plotted: green, measured time series; orange, mean stimulus-evoked response (estimated using deconvolution); purple, residual time series after removing the mean stimulus-evoked response. Horizontal black bars indicate the epoch of residual time series that was extracted for correlation analysis. **b**, Correlation coefficients for fearful face condition (left), neutral face condition (middle), and the difference in correlation (fearful − neutral), indicating the negative valence effect (right). Each square in “pink” colormap indicates the correlation between residual time series for a pair of ROIs under fearful face condition (left) or neutral face condition (middle). Each square in “hot” colormap indicates the difference in correlation between fearful and neutral conditions for a pair of ROIs (right). **c**, Correlation coefficients for happy face condition (left), neutral face condition (middle), and the difference in correlation (happy - neutral), indicating the positive valence effect (right). Each square in “pink” colormap indicates the correlation between residual time series for a pair of ROIs under happy face condition (left) or neutral face condition (middle). Each square in “hot” colormap indicates the difference in correlation between happy and neutral conditions for a pair of ROIs (right). **b-c** (left-middle), All squares in the fearful, happy, and neutral conditions showed correlation values significantly above 0 (*P* < 0.05, Wilcoxon signed rank test, Bonferroni-corrected for number of ROIs). **b-c** (right), Asterisks represent ROI pairs showing a statistically significant difference in correlation (\**P* < 0.05, One-sample t test, Bonferroni-corrected for number of ROIs). Number of unique participants scanned at 3T BOLD and 7T BOLD who were also scanned in the face localizer experiment: *n*=15 (see Table 1).

We found that all visual areas are highly and positively correlated with one another during viewing of fearful, happy, and neutral faces (Fig. 2b-c, left and middle, Supplementary Tables 1-3). It is important to note that these strong correlations were not a result of stimulus-evoked responses, since they were regressed out of the measured time series. Instead, these strong correlations likely reflect connectivity among visual cortical areas^17^. We also observed significant positive correlations between the amygdala and the rest of visual cortex (Fig. 2b-c, left and middle, 1^st^ column), though amygdala-cortical correlations were substantially lower than cortico-cortical correlations. The lower amygdala-cortical correlations could be due to considerably smaller response amplitude in the amygdala (relatively to visual cortex), which has been observed in both human and monkey studies^29,30^. The generally small amygdala-cortical correlations could also reflect signal contamination from a nearby vein^31^, large physiological noise in the amygdala, or a combination of both factors. Finally, the correlation differences between fearful and neutral face conditions (i.e., the negative valence effect) were evident between the amygdala and almost all visual cortical areas (Fig. 2b, right, 1^st^ column), consistent with findings of diffuse feedback-like projections from basal amygdala to a number of visual areas, including V1 in monkeys^15,32^. In contrast, inter-area correlation valence effect was not evident between V1 and any other cortical area, including V2 or V3 (Fig. 2b-c, right, 2^nd^-3^rd^ columns), suggesting that intracortical feedback is unlikely the source of the valence effect in V1.

Next, we tested the retinotopic specificity of the amygdala-V1 inter-area correlation valence effect. Functional imaging, brain stimulation and behavioral results suggest that feedback from ventral cortical areas projects to the foveal confluence of early visual cortex^18–20^. In contrast, anatomical projections from the amygdala to V1 are retinotopically diffuse, and project widely throughout V1^15,32^. Hence, if the valence effect in V1 were due to communication with other visual cortical areas, we would expect to observe enhanced correlations only at the fovea. In contrast, if it is due to feedback from the amygdala, we would expect diffuse correlation enhancements, evident at both the fovea and periphery. We constructed a peripheral V1 ROI, extending from beyond the stimulus representation all the way out to 88 deg of visual angle. We observed robust correlation enhancements between the amygdala and peripheral V1 (Fig. 2b-c, right), consistent with diffuse feedback projections from basal amygdala to V1.

### Layer-specific valence effect in V1

To determine the anatomic source of valence information in V1 (Fig. 3a), we used high field strength fMRI at 7T combined with VASO^23,24^ scanning. By measuring CBV responses across cortical layers (Fig. 3c-e), our approach enabled layer-specific measurements of both feedforward and feedback activity in V1 and minimized confounds introduced by draining veins that are inherent to BOLD fMRI^33^. Although VASO measurements typically have lower signal-to-noise ratios than BOLD, they are less contaminated by high-amplitude responses in superficial layers due to large draining veins^34^ (Fig. 3c). Finally, we note that VASO responses have the opposite sign from BOLD responses, as was evident in the 180° shift in the response phase, indicating that the VASO responses reached a minimum at roughly the same point in time in which BOLD responses reached their maximum (Fig. 3d). This observation indicates that the VASO pulse sequence that we used was indeed sensitive to CBV, rather than residual BOLD effects^23^, which would be expected to share the same response phase.

**Fig. 3:**
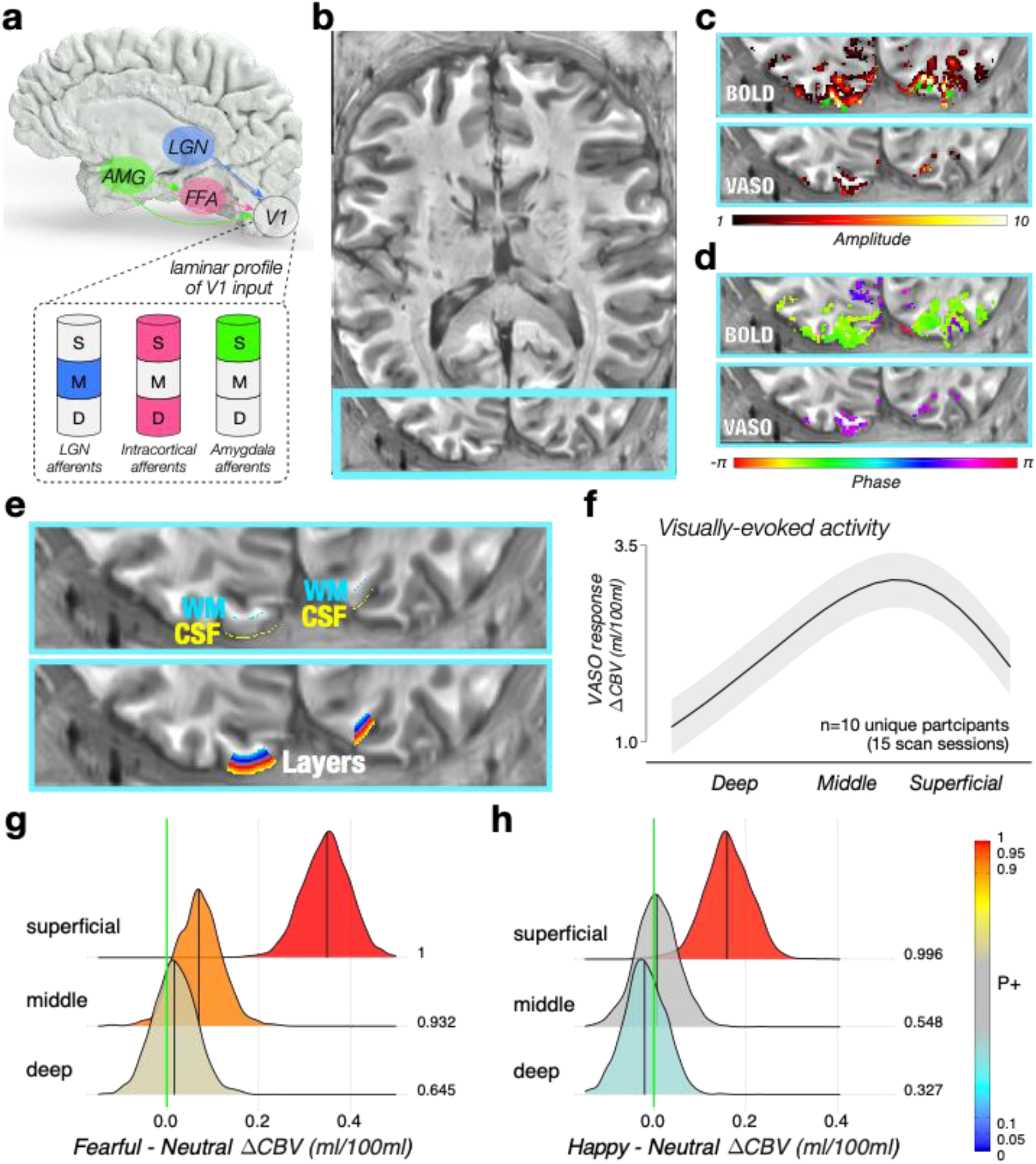
Facial valence modulation specific to superficial layers of V1. **a**, Three input pathways to V1 have distinct laminar profiles: LGN afferents terminate in the middle layer^36,37^, cortico-cortical afferents, such as from FFA, terminate in the superficial and deep layers^15^, and amygdala afferents terminate exclusively in the superficial layer^15,16^. AMG: amygdala, FFA: fusiform face area, LGN: lateral geniculate nucleus, D: deep layer, M: middle layer, S: superficial layer. **b**, Axial slice of a T1-weighted anatomical image generated from VASO timeseries^23^. Light blue line corresponds to the field of view shown in panels c-e. **c**, Response amplitude to face stimuli measured with BOLD (top) and VASO (bottom). VASO measurements are lower in amplitude but are more closely colocalized with cortical gray matter. Green arrows in the BOLD image (top) indicate high-amplitude responses in veins. **d**, Phase (timing) of the best-fitting sinusoid. BOLD (top) and VASO (bottom) are known to have opposite signed responses^23^, as indicated by the 180 deg shift in response phase. **e**, The central V1 ROI was defined in each participant based on retinotopic analysis^38^ and further constrained by demarcating the white matter (WM; cyan) and CSF (yellow) boundaries (top). Between these WM and CSF boundaries, 21 cortical depths were generated with LAYNII software^39^ (bottom). **f**, Percent change in VASO (in units of ml per 100 ml CBV) to all faces (pooled across fearful, neutral, and happy expressions) as a function of relative cortical depth between WM (left) and CSF (right). The black line shows the fitted average cross-layer profile among the three conditions, while the shaded band indicate the uncertainty range of one standard error. **g**, Posterior distribution of fearful – neutral VASO responses (in units of ml per 100 ml CBV) as a function of cortical depth. **h**, Posterior distribution of happy – neutral VASO responses (in units of ml per 100 ml CBV) as a function of cortical depth. **g-h**, Hue indicates the strength of statistical evidence according to the BML model^27^, shown through *P*+ (value at right side of each posterior distribution), the posterior probability of the valence effect at each cortical depth’s being positive conditioning on the adopted BML model and the current data. The vertical green line indicates zero effect. Cortical depths with strong evidence of the valence effect can be identified as the extent of the green line being farther into the tail of the posterior distribution. **f-g**, Number of unique participants scanned at 7T VASO: n=10 (15 scan sessions, see Table 1).

Depth dependent measurements of CBV (using VASO) in response to face stimuli exhibited two important characteristics (Fig. 3f-h). First, we observed a single peak in the mid-cortical depths of V1 for voxels corresponding to the retinotopic location of the stimuli (Fig. 3f). This peak was evident for each of the facial expressions (Supplementary Fig. 5) and was likely related to stimulus-evoked activity intrinsic to V1^35^. Second, the difference in response amplitude between fearful and neutral faces, and between happy and neutral faces, was most pronounced in superficial cortical depths of V1 (Fig. 3g-h). This laminar profile of the valence effect is consistent with direct projections from the basal amygdala, which terminate exclusively in the superficial layers of V1^15,16^.

The laminar profile of CBV in central V1 is consistent with two distinct sources of activity: stimulus-related drive, both from the LGN in middle cortical layers and recurrent local connections^35^ (Fig. 3a, blue region/layer) and direct afferents from the amygdala to superficial cortical layers^15^ (Fig. 3a, green region/layer), and is inconsistent with feedback from downstream cortical areas (Fig. 3a, magenta region/layer). This profile was highly reproducible across scan sessions on different days within participants (Supplementary Fig. 6). Moreover, the laminar specificity of the valence effect in V1 reported here is in line with the termination pattern of amygdala projections at the border between cytoarchitecturally defined layers I-II in V1 of the macaque monkey^15^, suggesting that the valence effect in V1 may be accomplished through direct projections from the amygdala, rather than feedback from other cortical areas, such as the FFA.

### Retinotopically-diffuse valence effect in V1

One implication of the inter-area correlation analysis (see above) is that feedback from the amygdala is retinotopically diffuse, and not restricted to the stimulated region of visual cortex. To directly test this hypothesis, we examined the retinotopic specificity of response amplitude modulation with facial expression in V1. We discovered that the valence effect in V1 (Fig. 1b-c, Supplemental Figs. 1-3) was not confined to the retinotopic location of the centrally presented face stimuli (Fig. 4a-b). Instead, it was present throughout V1, extending beyond the retinotopic representation of the stimulus, and even beyond the boundary of the stimulus display (Fig. 4c).

**Fig. 4:**
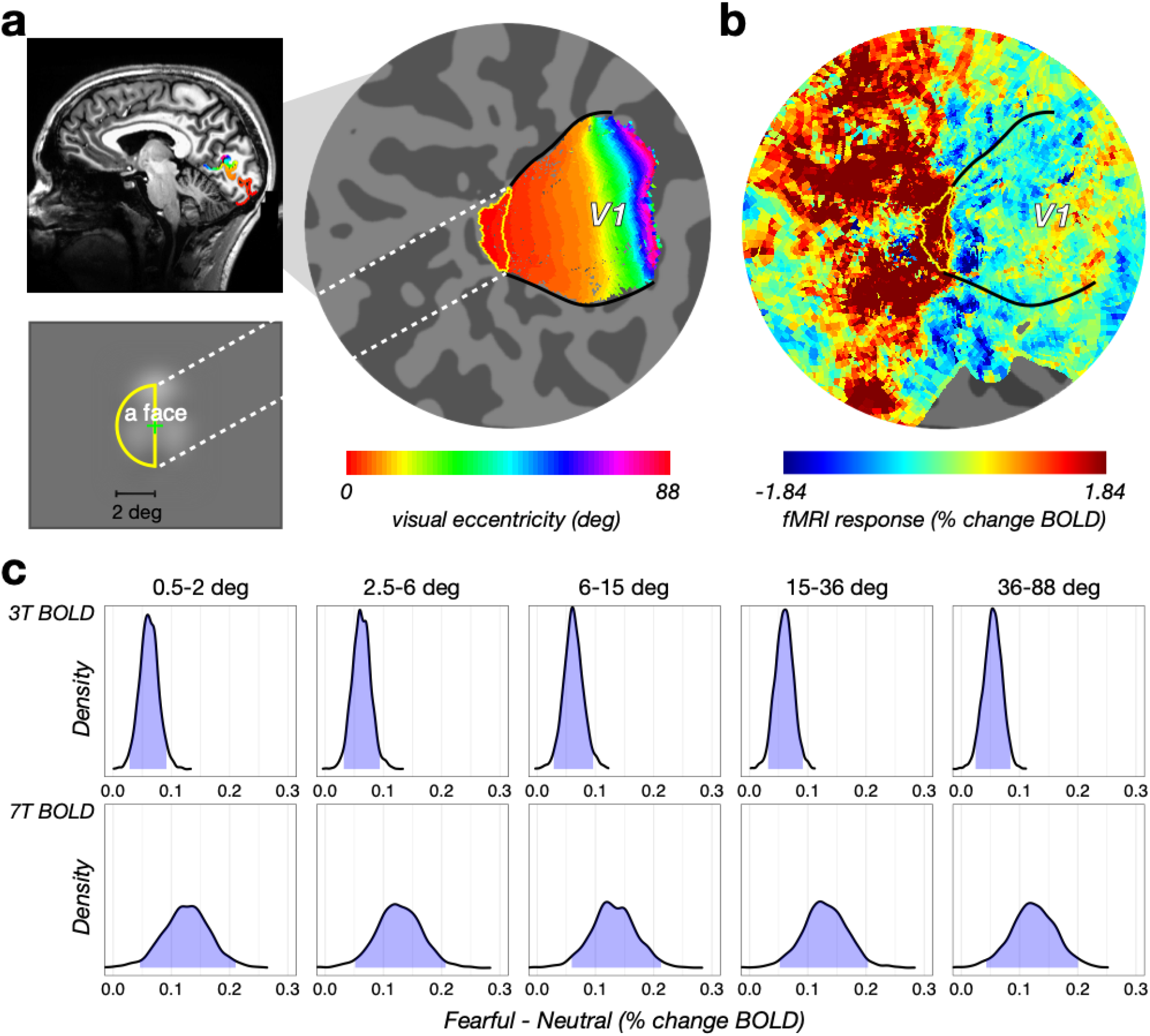
Facial valence modulation as a function of visual eccentricity. **a**, Face stimuli subsumed a 4 deg x 6 deg ellipse centered at the fovea and were expected to evoke responses in a retinotopically-identified region of V1, shown in a mid-sagittal slice (top left) and on a computationally flattened patch of early visual cortex (right). Hue indicates visual eccentricity. Yellow contour on the flat map indicates the retinotopic location of the face stimulus (bottom left). **b**, Visually-evoked BOLD response to all faces (same participant as in panel a). Black curves indicate V1/V2 boundary. Spatial pattern of visual response exhibits a strong positive response at the retinotopic location of the stimulus (red voxels), a surrounding negative penumbra at mid-eccentricities (dark blue voxels), and a return to baseline at far eccentricities (cyan, green, and yellow voxels). **c**, Valence modulation evident at all visual eccentricities. The statistical evidence for the elevated activity in response to fearful relative to neutral faces was substantial at all visual eccentricities. Under the posterior distribution of each eccentricity bin, the blue shadow indicates the 95% uncertainty intervals of the valence effect (fearful – neutral) with 5 eccentricity bins, separately for 3T BOLD (top) and 7T BOLD (bottom) scans. Number of participants scanned at 3T BOLD: *n*=14; number of participants scanned at 7T BOLD: *n*=14 (see Table 1).

We quantified fMRI response amplitude to fearful and neutral faces as a function of visual eccentricity. For each of the 20 participants in the 7T BOLD and 3T BOLD experiments, we performed a retinotopic analysis^38^ (Fig. 4a) to segment V1 into five iso-eccentricity bins (see Methods). The visual response averaged across different facial expressions exhibited a “half-Mexican hat” profile^40^ from central to peripheral V1 (Fig. 4b): a positive response at the fovea and extending out to 2 deg (dark red, radius: 0.5 – 2 deg), corresponding to the retinotopic location of the foveally presented faces (Fig. 4a, bottom), surrounded by a negative response penumbra (dark blue, radius: 2.5 – 6 deg), and then a return to baseline at more eccentric portions in V1 (green and yellow, radius: 6 – 88 deg). We found that the valence effect was evident at all eccentricities, and its magnitude did not differ as a function of eccentricity (Fig. 4c). Thus, the valence effect in V1 may be characterized as an additive effect riding on top of the stimulus-evoked response^41^, rather than a multiplicative gain modulation, as has been observed with spatial attention^42^. Although recent evidence suggests that amygdala neurons represent the spatial location of emotionally significant stimuli^43,44^, the valence effect observed in peripheral portions of V1 in the absence of visual stimulation is consistent with the observations from anterograde tracer studies that amygdala afferents are diffusely distributed throughout V1 in macaque^15,32^.

### Behavioral performance during scanning

Participants performed a gender judgment task on the face stimuli, orthogonal to facial expression, in order to help ensure a constant attentional state (but see^45^). There was no significant difference in gender judgment accuracy across the three fMRI experiments (3T BOLD: 92.07±3.90%, 7T BOLD: 91.66±3.31%, 7T VASO: 92.35±4.71%, one-way ANOVA: *F*(2,35) = 0.094, *P* = 0.910, Supplementary Fig. 7a). To examine potential within-subject performance differences across facial expressions, we collapsed performance within each of those who participated in multiple scan sessions (see Table 1). There was a significant main effect of facial expression on gender judgment performance, consistent across accuracy, reaction time (RT, correct trials only), and inverse efficiency score (IES) measures (one-way repeated measures ANOVA: all *P* values < 0.001). Specifically, performance when fearful faces were presented (accuracy: 89.75±4.18%, RT: 641±46ms, IES: 715±56ms) was significantly worse than performance when neutral (accuracy: 92.83±3.82%, RT: 637±42ms, IES: 687±53ms) or happy faces (accuracy: 94.53±3.03%, RT: 631±43ms, IES: 668±47ms) were presented (Supplementary Fig. 7b-d). Importantly, test-retest reliability of task performance showed that behavior was highly consistent across scan sessions on different days within participants (Supplementary Fig. 8).

## Discussion

Studying the neural circuits underlying emotion processing provides a unique window into how primate brains evolved to deal with the challenges of living in large social groups. In this study, we showed that the facial valence effect is widely distributed, present in most cortical areas that respond to face stimuli. We used high-resolution layer-specific fMRI to characterize the valence effect in V1. We found that the valence effect was limited to superficial layers but was not limited to the retinotopic location of the face stimuli. These two observations are consistent with the known anatomical connectivity between the amygdala and V1^15,32^ and suggest that the valence effect in V1 arises through this direct pathway rather than through indirect pathways involving feedback from other cortical areas. Our neuroimaging results in V1 are consistent with behavioral observations that emotion affects early visual processing^46^, highlighting the role of the amygdala in enhancing the sensory processing of emotionally-salient stimuli.

A large number of prior fMRI studies have reported facial valence effects^5,47,48^, and these studies have generally focused on a limited set of cortical and subcortical peak activation loci that are thought to subserve the processing of emotional facial expressions. We measured BOLD activity throughout the entire brain and found that the valence effect was present in nearly all cortical areas that exhibited responses to face stimuli (Fig. 1b-e). We make three points. First, the impact of facial expression may be more widespread than is commonly appreciated, involving brain areas that are not classically considered part of an emotion-processing network. Second, while we focused our analysis on areas that could be reliably identified with a retinotopic atlas^38,49^, we observed valence effects in a number of high-level cortical areas that are not part of the retinotopic atlas, including the superior temporal sulcus, the superior parietal lobule, and the inferior prefrontal sulcus. Third, despite converging evidence of valence sensitivity in early visual cortex from human EEG/ERP studies^50,51^, recordings in awake monkey^52^, and computational modeling^53^, fMRI evidence for valence sensitivity in human early visual cortex has been conflicting, with clear effects reported in studies using face stimuli^5,54^, and studies using emotional scene and applying decoding analysis^22,53^, but not studies using emotional scene and applying univariate analysis^55–57^. Our results demonstrate clear and reliable valence sensitivity throughout human visual cortex, including in V1.

It is likely that there are multiple mechanisms involved in processing affective stimuli (e.g., changes in perceptual processing, arousal, memory, and motor output). In particular, it is conceivable that eye movement patterns associated with viewing neutral and fearful faces could have interacted with our results, though we think neither is very likely. First, while participants were instructed to fixate throughout the entire experiment, microsaccades were inevitable. It is possible that microsaccade rate and/or direction were modulated by facial valence, as with spatial attention^58^, but it is difficult to see how changes in microsaccades could have produced the pattern of activity that we observed. Each microsaccade would cause some degree of retinal stimulation when stable visible features (e.g., the stimulus or the edge of the screen) move across the retina. However, we found that the valence effect extended from 0.5 deg all the way to 88 deg, well beyond both the stimulus and the screen edge (Fig. 4). Moreover, microsaccades would be expected to evoke positive BOLD responses in visual cortex^59^, but we found negative responses beyond the stimulus representation, most likely due to surround suppression (Fig. 4). Second, it is conceivable that fearful faces caused pupil dilation, which would in turn, allow more photons to enter the eye, resulting in a global response in visual cortex. However, the percentage change in pupil size needed to effect such a large change in cortical activity would need to be dramatic^60^.

Many layer-specific fMRI studies have characterized feedback responses in the absence of bottom-up, feedforward drive^54,61,62^. The layer profile that we measured contained both feedforward signals arising from the LGN, as well as feedback signals related to facial valence. Feedforward responses to flickering checkerboard stimuli have been characterized in a recent layer-specific fMRI study^63^, in which the largest fMRI response was observed in a middle-deep cortical depth that colocalizes with the stria of Gennari^64^, a band of heavily-myelinated fibers within layer 4B containing synapses from geniculocortical projection. MRI images of the stria of Gennari can be obtained with a variety of MR contrast mechanisms (for a review, see Ref^65^). While we did not acquire a scan enabling us to identify the stria of Gennari in our study, we note that the peak response across face stimuli was evident in more superficial cortical depth than the expected depth of the stria of Gennari. One possible explanation for this depth profile is related to the widely-characterized superficial bias from draining veins^66^. However, this explanation is unlikely to account for our results because the layer profile that we measured decreased at the most superficial cortical depths where the vascular effects are expected to be strongest. Alternatively, the cortical depth profile that we measured with VASO matches the laminar profile of the local field potential in macaque V1 in response to grating stimuli, which is largest in layers 2/3 and 4B^35^. Activity in these laminae is thought to reflect stimulus-induced local recurrent activity, which may be a more pronounced source of net neural activity than the feedforward drive from the LGN to layer 4C.

We observed a facial valence effect only in the superficial layers of V1, and we interpret this as evidence for feedback to these layers. One alternative explanation for this depth-dependent response profile is related to the widely-characterized superficial bias from draining veins, in which the largest response amplitudes are observed in the superficial layers^66^. Even though VASO is thought to mitigate the impact of draining veins^23,24^, it is conceivable that BOLD contrast contaminates the VASO measurement to some degree. However, we think this is unlikely for two reasons. First, the VASO responses in our experiment were signal decreases, i.e., negative responses (Fig. 3, but note that the responses were multiplied by −1). This suggests that the removal of the BOLD component of the signal was successful. Second, the layer profile that we report (Fig. 3f) exhibited a clear and prominent decrease at the most superficial cortical depths, rather than a linear increase toward the pial surface as would be expected from a BOLD layer profile. This observation suggests that the activity profile reflects changes in CBV rather than a vascular confound.

Feedforward, stimulus-driven patterns of activity have been studied extensively and in great detail in human visual cortex using fMRI (for reviews, see Ref ^67^). By contrast, relatively little is known about the role of feedback in human visual cortex. This is mainly because studies of feedback have been limited to invasive measurement methods^68–70^, and hence are beyond the purview of fMRI and other noninvasive methods of measuring cortical activity in humans. However, the rapidly-expanding field of high-resolution fMRI has begun to elucidate the crucial role of feedback in shaping visual responses^61,62^. The majority of studies in V1 have focused on the role of cortico-cortical feedback; comparatively, little is known about other feedback projections to V1. Here, we applied layer-specific fMRI to understand how visual cortical responses are modulated by fearful faces, and in particular, the role the amygdala plays in this process. Note that amygdala activation may not be specific to fear^71^ nor to facial expressions^72^. The amygdala responds to a variety of biologically relevant stimuli, such as animate entities^73^, ambiguous or unpredictable cues^74^, and social category groups^75^.

In addition to the feedforward response we measured, the neural pattern of valence modulation we characterized—functionally correlated between the amygdala and both central and peripheral V1 (Fig. 2), specific to the superficial cortical depths of V1 (Fig. 3), retinotopically non-specific, and evident throughout the spatial extent of V1 (Fig. 4)—suggests that sensitivity to facial valence in V1 arises from direct anatomical projections from the amygdala. This pattern is inconsistent with the alternative anatomical pathway we considered in the introduction. That is, valence information computed in the amygdala reaches V1 via cortico-cortical feedback projections from extrastriate areas^22^. Although many visual areas exhibited a valence effect (Fig. 1b-c) and also send feedback projections to V1^76^, projections from these areas terminate in both superficial and deep layers^77^, inconsistent with the layer profile we observed. The layer-specific and retinotopically non-specific pattern is further inconsistent with two additional alternative pathways we consider. One alternative pathway is the cholinergic projections from the basal forebrain. The basal forebrain receives prominent inputs from the amygdala^78^ and also sends projections to V1^79^. However, afferents from basal forebrain to V1 terminate in all layers and are most dense in layers 1, 4 and 6 in macaque^80^, making this pathway an unlikely candidate to explain our fMRI results. The other possibility is that the valence information is computed in the pulvinar^54^, not in the amygdala, and this information is then transmitted to V1. Pulvinar afferents are mainly located in layer I of V1 in primates^81^, consistent with our layer fMRI results. However, these pulvinar-V1 projections are retinotopically specific^82^ and would not produce the diffuse pattern of valence modulation that we observed. We, therefore, conclude that direct projections from the amygdala are the most likely source of valence modulation in V1.

Our fMRI experiment employed a block design with three different facial expressions (happy, neutral, fearful) with interleaved fixation blocks that were shorter (half the duration) than the face blocks. With the relatively short fixations block, the post-stimulus undershoot from one face block overlapped with the beginning of the response in the next block of trials (Supplementary methods; Supplementary Fig. 9). This design is derived from classic experiments in which interleaved fixation blocks were shorter than stimulation blocks (i.e., 30 s stimulus blocks interleaved with 20 s fixation blocks in Kanwisher et al., 1997^83^; 9s stimulation blocks interleaved with 6s blank screen in Levy et al., 2001^84^ and Hasson et al., 2002^85^). The fMRI BOLD response approximates a shift-invariant linear system^86–88^, which makes it possible to deconvolve overlapping responses from different conditions, provided the time series is sufficiently long and the conditions sufficiently randomized and counter-balanced^89^.

There are two important assumptions when applying this design to layer fMRI. The first is that the linearity of the response applies to measurements at each cortical layer. For example, it is conceivable that response at one layer conforms to the linearity assumptions, but responses at other layers deviate from linearity to some degree, perhaps due to directional blood pooling towards the pial surface. Initial studies suggest that linearity assumptions do apply to layer-specific fMRI^90,91^, but this issue does deserve greater attention. The second assumption is that the VASO measurements are linear in the same way as BOLD measurements. VASO fMRI is an indirect measurement of CBV, which is thought to exhibit linearity^92^. However, more work on the linearity of VASO is warranted. Nonetheless, slight deviations from linearity, if present in our measurements, are unlikely to account for the results that we report here.

Coregistration between anatomical and functional data is a major challenge for high-resolution fMRI^93^. We overcame this challenge by adopting an approach that did not require coregistration. Specifically, we used a distortion-matched T1 weighted anatomical volume^94^ that was computed directly from the VASO measurements. We then hand segmented the cortical ribbon of central V1 in the native space of data acquisition.

It is well established that emotional faces receive more attention than neutral stimuli^46,95^. In our study, participants performed a facial gender judgment task, orthogonal to emotional expression in order to decrease potential attentional differences between conditions. Nevertheless, we observed behavioral differences in gender judgment between emotional expressions. One potential reason for this behavioral effect could be related to subtle differences in low-level image features across expressions. We did control for low-level image statistics using the SHINE toolbox^96^, and there were no global differences in image luminance, contrast, or spatial frequency across all three expressions. However, we cannot rule out the possibility of local differences in image statistics between facial expressions, which are not normalized by the SHINE toolbox. A second possibility is that gender may be less discriminable in fearful faces compared to neutral or happy faces, which led to worse gender judgment performance in the fearful condition. However, as evident from a pixel-level representational similarity analysis (RSA), the largest representational distance between female and male faces was found in the fearful condition, suggesting that gender judgements should be more accurate in the fearful condition, which is the opposite of what we found. Finally, our behavioral results are most consistent with difficulty in disengaging attention from faces with negative valence (e.g., angry or fearful) relative to faces with positive valence (e.g., happy) or neutral faces^97^. Consistent with this third possibility, we observed behavioral performance worst for fearful, best for happy, with neutral faces intermediate between the two. Moreover, this pattern was qualitatively similar to that reported in another study using gender judgment of emotional faces^21^. Given the pattern of fMRI valence modulation (largest responses to fearful and happy faces, smallest for neutral faces), it is highly unlikely that the behavioral difference in gender judgment across facial expressions could have given rise to the pattern of fMRI results reported here. Thus, the valence-specific effect we observed in fMRI was not simply due to differences in task difficulty.

The facial valence effect in retinotopic visual cortex we found are broadly consistent with a recent EEG-fMRI study that demonstrated affective scene decoding in retinotopic visual cortex^22^. In that study, however, the amplitude of the late positive potential (LPP)—an index of reentrant processing from the amygdala back to visual cortex^98^—correlated only with the fMRI decoding accuracy in ventral visual cortex, but not in early or dorsal visual cortex, suggesting that the valence effect in early visual cortex may arise from reentrant signals propagated to V1 from ventral visual cortex. This may suggest that valence-related feedback signals are stimulus specific, with face stimuli and perhaps animate objects more generally^73^, engaging the circuitry from basal amygdala to V1, and scene stimuli engaging connectivity between ventral visual cortex and V1. Regardless of stimulus type, the valence effect occurs throughout visual cortex in both studies. It is known that face and scene stimuli are associated with distinct patterns of brain activity beyond the amygdala^99^. Two key factors may underlie potentially distinct mechanisms of emotional face and scene processing. First, the heterogeneity in image statistics is smaller across faces than across natural scenes. Second, compared to the direct communicative role of facial expressions, the emotional and social aspects of scene processing are commonly perceived as more indirect and secondary. Thus, future work will need to use network analysis of whole brain dynamics across different imaging modalities to determine whether these widespread valence effects are due to direct influence from the amygdala, feedforward inputs from V1, or a combination of both.

## Methods

### Participants

A total of 53 2-hour scan sessions from 34 healthy right-handed volunteers (age 21-42 years, 16 females) from the DC/MD/VA tri-state area were collected in this series of experiments (7T VASO, 7T BOLD, and 3T BOLD). Each volunteer participated in 1-4 scanning sessions across three experiments. All participants granted informed consent under an NIH Institutional Review Board approved protocol (93-M-0170, ClinicalTrials.gov identifier: NCT00001360). Two 7T VASO participants were scanned with personalized headcase (Caseforge, https://caseforge.co) to reduce head motion and a separate consent was obtained prior to the headcase scanning appointment.

Based on conservative head motion parameter estimates across different magnetic strength or voxel size, seven 7T VASO scan sessions from six participants were excluded due to excessive head motion (>1 mm translation or >1° rotation within each run and/or >2 mm translation or >2° rotation across runs within a single scan session). Data from an additional 3 participants were further excluded because of technical errors, lack of scan time, or outlier behavioral performance (>3 SD below mean accuracy). Hence, the final dataset reported here includes a total of 43 scan sessions from 25 participants (average age 25.9 years, 12 females, see Table 1), consisting of 14 scan sessions from 14 unique participants at 3T BOLD, 14 scan sessions from 14 unique participants at 7T BOLD, and 15 scan sessions from 10 unique participants at 7T VASO, among whom 5 were scanned twice to evaluate test-retest reliability of VASO (see Supplementary Fig. 6).

### Visual stimuli

Participants viewed face stimuli that varied in emotional expression while performing an orthogonal gender judgment task. The stimuli consisted of 168 facial identities from 56 unique individuals (28 female and 28 male) images of faces taken from the Emotion Lab at the Karolinska Institute (KDEF)^100^ and the NimStim database^101^. All face stimuli were preprocessed using the SHINE toolbox^96^ to control for low-level image statistics. There was no global luminance difference across expressions (one-way ANOVA: *F*(2,165) = 0.138, *P* = 0.871), no effect of contrast on expression (one-way ANOVA: *F*(2,165) = 0.041, *P* = 0.960), and no effect of spatial frequency on expression, *F*(2,165) = 1.04, *P* = 0.3535).

An RSA on pixel-level discriminability between female and male faces in each expression group revealed a significant effect of expression on gender discriminability (one-way ANOVA: all *F* values > 148.03, all *P* values < 0.001 across Euclidian distance, correlation distance and cosine distance). Specifically, facial gender in the fearful condition (Euclidian distance: 4987±25) was higher in discriminability than that in the neutral (Euclidian distance: 4405±25, independent samples t-test: *t*(1566) = 16.5, *P* < 0.001) or happy condition (Euclidian distance: 4872±25, *t*(1566) = 3.26, *P* < 0.001). Moreover, facial gender in the happy condition was higher in discriminability than that in the neutral condition (independent samples t-test: *t*(1566) = 13.149, *P* < 0.001). The size of the emotional face stimuli was also matched across 3T and 7T scans: all faces with different emotional expressions extended 4 deg horizontal and 6 deg vertical. Participants fixated a small (1 deg) green fixation cross for the duration of each run.

The localizer scan contained 104 images in each of the three categories: faces, objects, and scrambled objects (adapted from Refs. ^102,103^). Different from the emotional faces shown in the gender judgment task, the face images used in the independent face localizer were obtained from the Face Place database (http://www.tarrlab.org). Prior to the first scanning session, all participants practiced both the gender judgment task and the one-back task (face localizer), if included in the scan session, for several minutes.

All tasks were run using MATLAB 2016b (MathWorks, MA) and MGL toolbox^104^ on a Macintosh computer. Stimuli were displayed on a 32″ 1920×1080 MRI-compatible LCD screen (BOLDscreen 32 LCD for fMRI, Cambridge Research Systems) at the head end of the bore in 3T and were projected onto a rear-projection screen using a 1920×1080 LED projector (PROPixx, VPixx Technologies Inc) at 7T. In all experiments, stimulus presentation was synchronized with the fMRI scanner on each TR.

### Gender judgment task

In both 3T BOLD and 7T BOLD experiments, each run consisted of three repeats of each facial expression condition (fearful, neutral, happy) and ten repeats of the fixation block. Within each facial expression block, each face was presented for 900 ms with a 100 ms interstimulus interval (ISI) while a green fixation cross remained at the center of the screen at all times. Each fixation block lasted 10 s and each face block lasted 20 s in the 3T BOLD experiment. Hence, each run lasted 4 min 40 s in total. Similarly, each fixation block lasted 9 s and each face block lasted 18 s in the 7T BOLD experiment; thus each run lasted 4 min 12 s in total.

In the 7T VASO experiment, each run consisted of six repeats of each facial expression condition (fearful, neutral, happy) and 19 repeats of the fixation blocks. There were 16 faces presented in each face block. Each face was shown for 1100 ms with a 106.25 ms ISI. Thus, each face block lasted 19.3 s and each fixation block lasted for 9.65 s, and each run lasted 8 min 53 s.

Participants performed a gender judgment task (press “1” for female, “2” for male) for each face stimulus, orthogonal to facial expressions. Feedback on task performance (percent correct) and real-time head motion estimates were given to the participant shortly after each run; no feedback was given during scanning. Participants were not told the purpose of the study but were debriefed following the last scan session upon request.

### Face localizer

An independent face localizer task^102,103^ was performed for all but three participants in the 3T BOLD experiment and all but four participants in the 7T BOLD experiment. The localizer was run using a block design with stimuli from three categories: faces, objects, and scrambled objects. The design and timing were matched to those in the gender judgment task, such that 1) each run consisted of three repeats of each category and ten repeats of the fixation block, and 2) within each category, the stimulus was presented for 900 ms with a 100 ms ISI while a green fixation cross remained at the center of the screen at all times. Like the gender judgment task, each face localizer run lasted 4 min 40s in total in the 3T BOLD experiment and 4 min 12s in the 7T BOLD experiment. Participants were instructed to indicate an immediately repeating image among 16 images per block (a one-back task: press “1” for same, “2” for different) and responses were made using the right index finger via a MR compatible button glove. This response instruction was designed to maximally engage participants while keeping the task relatively easy (performance was at ceiling, e.g., accuracy: 95.3±2.1%).

#### Image acquisition

3T BOLD fMRI data were collected on a Discovery MR750 scanner (GE Healthcare, Waukesha, WI, USA) with a 32-channel receive head coil, while 7T BOLD and 7T VASO fMRI data were acquired on a MAGNETOM 7T scanner (Siemens Healthineers, Erlangen, Germany) with a single-channel transmit/32-channel receive head coil (Nova Medical, Wilmington, MA, USA). Both 3T and 7T scanners were located at the functional magnetic imaging core facility on the NIH campus (Bethesda, MD, USA). For 7T scans specifically, a 3rd order B0-shimming was done with four iterations. The shim volume covered the entire imaging field of view (FOV) and was extended down to the circle of Willis to obtain sufficient B0 homogeneity for VASO inversion.

### BOLD scan parameters

3T BOLD fMRI data were acquired using multi-echo gradient-echo echo planar (EPI) sequence (TR = 2000 ms, TE1 = 12.5 ms, TE2 = 27.6 ms, TE3 = 42.7 ms, voxel size = 3.2 x 3.2 x 3.5 mm, flip angle = 75°, echo spacing = 0.4 ms, grid size = 64 x 64 voxels, 30 slices). 7T BOLD fMRI data were acquired using a gradient-echo EPI sequence (TR = 1500 ms, TE = 23 ms, voxel size = 1.2 x 1.2 x 1.2mm, flip angle = 55°, grid size = 160×160 voxels, 42 slices).

### VASO scan parameters

7T VASO data were acquired using an inversion recovery prepared 3D-EPI sequence, which was optimized for layer-specific fMRI in human visual cortex^23^. Parameters of inversion recovery preparation were as follows: The adiabatic VASO inversion pulse is based on the TR-FOCI pulse, with a duration of 10 ms and a bandwidth of 6.3 kHz. The inversion efficiency was adjusted by the implementation of a phase skip of 30 deg to minimize the risk of inflow of fresh non-inverted blood into the imaging region during the blood nulling time. 7T VASO data were acquired using a 3D-EPI readout with the following parameters: 0.82 x 0.82 x 0.82 mm, FOV read = 133 mm, 26 slices, whole k-space plane acquired after each shot, FOV in the first phase encoding direction = 133.3% of FOV in the readout direction, TE = 24 ms, GRAPPA 3, partial Fourier of 6/8. To account for the T1-decay during the 3D-EPI readout and potential related blurring along the segment direction, a variable flip angle (FA) was applied across segments, which started from 22° and then exponentially increased until reaching a desired flip angle of 90°.

The acquired time series consisted of interleaved BOLD and VASO images, with TRBOLD = 2737 ms and TRVASO = 2088 ms, resulting in effective TRVASO+BOLD = 4825 ms. A more detailed list of scan parameters used can be found: https://github.com/tinaliutong/sequence.

Imaging slice position and angle were adjusted individually for each 7T VASO participant so that the slice prescription was parallel to each participant’s calcarine sulcus (visualized on the sagittal plane prior to the scan, see Supplementary Fig. 6a). We also ran the retinotopic atlas analysis based on each participant’s T1-weighted MPRAGE MRI, acquired in a separate session prior to the main experimental scan session. This was used to guide slice prescription, aiming to maximally cover V1 in each participant. After slice prescription, a third order B0-shimming was done with four iterations. The shim volume was parallel to the slice prescription.

Image reconstruction was done in the vendor-provided platform (Siemens software identifier: IcePAT WIP 571) and was optimized with the following set-up to minimize image blurring and increase tSNR at high resolution. GRAPPA kernel fitting was done on FLASH autocalibration data with a 3×4 kernel, 48 reference lines, and regularization parameter χ = 0.1. Partial Fourier reconstruction was done with the projection onto convex sets (POCS) algorithm with eight iterations. Data of each coil channel were combined with the sum of squares.

### Structural MRI

Within the same 3T scan session, anatomical images were acquired in each individual for co-registration purposes using a 3D Magnetization-Prepared Rapid Acquisition Gradient Echo (MPRAGE) sequence with 1 mm isotropic voxels, 176 sagittal slices, acquisition matrix = 256 x 256, TI/TE/TR = 900/1.97/2300 ms, flip angle = 9, ° GRAPPA = 2, scan time = 5 min 21s. The 3T anatomy was also used for co-registration of all 3T BOLD participants and 8 of 14 7T BOLD participants (who participated in both 3T BOLD and 7T BOLD scans). In other 7T participants, a 0.7 mm isotropic resolution T1-maps were collected covering the entire brain using an MP2RAGE sequence with TI1/TI2/TR/TE = 800/2700/6000/3.02 ms, FA1/FA2 = 4°/5°, 224 sagittal slices, matrix size = 320 × 320, scan time = 10 min 8 s. Before the VASO scan, we made sure all participants had prior MPRAGE data available, which was used to estimate the slice angle of the VASO scan.

#### fMRI time series preprocessing

All preprocessing steps were implemented in MATLAB 2016b using a combination of mrTools^104^ and AFNI software package^105^. Standard preprocessing of the 3T multi-echo gradient echo EPI data utilized the AFNI software program afni_proc.py. Data from the first 4 TRs were removed to allow for T1 equilibration and to allow the hemodynamic response to reach a steady state. Advanced automatic denoising was achieved using multi-echo EPI imaging and analysis with spatial independent component analysis (ICA), or ME-ICA^106,107^. Preprocessing of 7T BOLD data included head movement compensation within and across runs, linearly detrended, and high-pass filtered (cutoff: 0.01 Hz) to remove low-frequency noise and drift. For 7T VASO data, all time frames were first split into blood-nulled and blood-not-nulled (BOLD) groups. Motion correction was performed separately for each group. The time frames from each group were upsampled in time via cubic interpolation, and the first and last two upsampled time frames in each group were removed from each run. Next, CBV-weighted VASO signals were calculated as blood-nulled divided by blood-not-nulled (BOLD) at each time frame to remove BOLD contamination^34^ and multiplied by −1 to convert negative responses to positive responses.

#### fMRI statistical analysis

A standard general linear model (GLM) analysis was performed in mrTools^104^. The regressor for each condition of interest (faces, objects, and scrambled objects in the face localizer task, or fearful, neutral, happy in the gender judgment task) was convolved with a canonical hemodynamic response function. The correlation coefficients between each pair of ROIs, for fearful and neutral conditions, were computed based on the residual time series (measured response time series - predicted response time series estimated using deconvolution^108^) (Fig. 2a) and their difference in correlation (fearful - neutral) was entered into Wilcoxon signed-rank test (Fig. 2b). The beta weights (in units of percent signal change) and t statistics for the fearful, happy, and neutral conditions were entered into Bayesian Multilevel (BML) modeling (Figs. 3-4; Supplementary Figs. 5-6).

#### Bayesian Multilevel modeling

In Fig. 1, a region-based analysis was performed through BML modeling^27,109^. Specifically, the approach was applied to fMRI response amplitude *Y_crs_* of the three conditions with the Student’s *T*-distribution in an integrative framework,

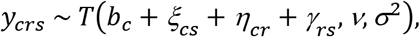

where *c*, *r*, and *s* index the 3 conditions, 15 ROIs, and 15 participants (Fig. 1 and Supplementary Fig. 4), respectively; *b*_*c*_ represents the effect of the *c*th condition at the population level; *ξ_cs_* codes the *s*th participant’s effect under the *c*th condition; *η_cr_* is the *r*th ROI’s effect under the *c*th condition; *γ_rs_* characterizes the *r*th ROI’s effect under the *c*th condition; *ν* and *σ*^2^ are the number of degrees of freedom and variances for the Student’s *T*-distribution whose adoption was intended to account for potential outliers and skewness. Three prior distributions were adopted as below,

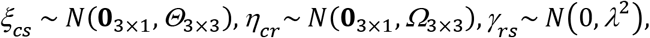

where *Θ*_3×3_ and *Ω*_3×3_ are 3 × 3 positive semidefinite matrices for the variance-covariance structures among the three conditions; *λ*^2^ is the variance for the interaction effects between regions and participants.

The BML model was numerically solved through the AFNI program RBA^30^ with 4 Markov chains each of which had 1,000 iterations. Noninformative hyperpriors were adopted for population-level effects *b*_*c*_; for the two variance-covariance matrices *Θ*_3×3_ and *Ω*_3×3_, the LKJ correlation prior was used with the shape parameter taking the value of 1 (i.e., jointly uniform over all correlation matrices of the respective dimension); a weakly-informative prior of Student’s half-*t*(3,0,1) was utilized for the standard deviation *λ*; the hyperprior for the degrees of freedom, *ν*, of the Student’s *T*-distribution was Gamma(2, 0.1); lastly, the standard deviation *σ* for the BML model was a half Cauchy prior with a scale parameter depending on the standard deviation of the input data. The consistency and full convergence of the Markov chains were confirmed through the split statistic *R̂* being less than 1.05. The effective sample size (or number of independent draws) from the posterior distributions based on Markov chain Monte Carlo simulations was more than 200 so that the quantile (or compatibility) intervals of the posterior distributions could be estimated with reasonable accuracy. The BML model’s performance was confirmed by the predictive accuracy through posterior predictive checks (Supplementary Fig. 4).

The BML modeling results show each region’s posterior distribution (Fig. 1c). Each contrast between two conditions *C* and *C* was expressed as a dimensionless modulation index 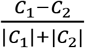, whose posterior distribution was represented through the posterior samples drawn from the Markov chain Monte Carlo simulations of the BML model. The strength of statistical evidence is shown through *P*+, the posterior probability of each region’s effect being positive conditioning on the adopted BML model and the current data. See the BML model performance in Supplementary Fig. 4.

The cross-layer profiles were fitted through smoothing splines (Fig. 3f, Supplementary Fig. 5, and Supplementary Fig. 6b). Specifically, we adopted thin plate splines as basis functions in a multilevel model to adaptively accommodate the nonlinearity of each cross-layer profile^110^. The measurement uncertainty (standard error) of the VASO response was incorporated as part of the input in the model, which was numerically solved through the R package mgcv^111^ to obtain the estimated cross-layer VASO profiles and their uncertainty bands.

In Figure 4, BML modeling was applied to fMRI response amplitude *Y_crs_* of the fearful and neutral conditions with the otherwise same framework as in Figure 1,

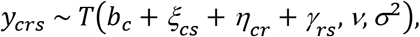

except where *c*, *r*, and *s* index the 2 conditions, 5 eccentricity bins in V1, and 14 participants each, in the 3T BOLD and 7T BOLD experiments, respectively (Fig. 4c).

#### Functional ROI definition

To examine the valence effect in a wide range of visual or face-selective areas (Fig. 1b), the amygdala and the FFA were functionally defined from the independent face localizer using a conjunction between t map of faces-objects (whole brain FDR<0.05) and R2 map (R2>0.1 for 11 scan sessions at 3T BOLD or R2>0.05 for 12 scan sessions at 7T BOLD). Based on a probabilistic atlas^25^, 25 visual areas per hemisphere were defined in these 23 scan sessions (from 15 unique participants). Next, visual areas with the same area label were combined across hemispheres (with IPS1-5, LO1-LO2, PHC1-PHC2, TO1-TO2, V1d-V1v, V2d-V2v, V3A-V3B, V3d-V3v, VO1-VO2 combined) and were further thresholded by R2 value in the independent face localizer (R2>0.1 at both 3T BOLD and 7T BOLD). For each participant, we also performed a retinotopic analysis using a probabilistic atlas^38^. The eccentricity map was visualized on a flat patch of early visual cortex and a portion of central V1 corresponding to the size and position of the face stimuli was highlighted by the yellow contour (Fig. 4a-b).

#### Inter-area correlation analysis

The goal of this analysis was to quantify the strength of correlations between brain areas using the component of the time series that was not driven by the task or the stimulus. To remove the stimulus-related component of the BOLD time series, we computed the residual time series after removing the mean stimulus-evoked responses. Mean stimulus-evoked responses were estimated using deconvolution^108^, separately for each ROI, in each scan session (see Fig. 2a for one run from an example participant). Specifically, a predicted time series *̂* was computed by multiplying the design matrix by the parameter estimates *x̂*. Next, the residual time series was computed by subtracting the predicted response time series from the measured response time series, *r* = *y* - *ŷ* Epochs of residual time series (each face block and its following fixation block) corresponding to each facial expression condition (fearful, neutral, happy) were concatenated across runs within a scan session and extracted for the inter-area correlation analysis.

Correlation coefficients between each pair of ROIs (defined above) were computed from the residual time series in each ROI corresponding to each facial expressions condition. The differences (fearful − neutral) in correlations were also computed (Fig. 2b). For participants who were scanned in multiple sessions, correlation coefficients were averaged between sessions.

#### VASO anatomy

To ensure the most accurate definition of cortical depths, we used the functional VASO data directly to generate an anatomical reference, termed VASO anatomy. It was computed by dividing the inverse signal variability across blood-nulled and blood-not-nulled images by mean signals. This measure is also called T1-EPI^23^, which provides a good contrast between white matter (WM), gray matter (GM) and cerebral spinal fluid (CSF; see Fig. 3b) in native EPI space.

#### Layering methods

All layer analyses were conducted in VASO EPI space. The VASO anatomy was first spatially upsampled by a factor of 4 in the in-plane voxel dimensions (X and Y directions) to avoid singularities at the edges in angular voxel space, such that the cortical layers can be defined on a finer grid than the original EPI resolution. We then coregistered each participant’s eccentricity map from the retinotopic atlas to the VASO anatomy from that particular session in order to generate an anatomical reference of central V1 in the native space of the data acquisition. This procedure ensured that no spatial resampling or loss of resolution (i.e., blurring) occurred in the functional EPI data. Cortical layers in V1 were defined on the z plane (axial slice) in reference to the borders between layer I of the GM and CSF, and between layer VI of the GM and WM. Across 26 slices in the Z direction, we first identified the slice with the highest R2 value (i.e., visually evoked response) within bilateral central V1. Next, we estimated twenty-one cortical depths between the two boundaries (Fig. 3e-f) using the LN_GROW_LAYERS program in the LayNii software^39^ (https://github.com/layerfMRI/LAYNII). The number 21 was chosen to enable more layers than the independent points across the thickness of the cortex, which can improve layer profile visualization and minimize partial volume effect between neighboring voxels. Note that we do not assume that these 21 layers are statistically independent measurements. We repeated the previous step for the slice above and below, and averaged the percent change in CBV signals across the 3 slices per layer. The number of voxels per layer in the upsampled resolution in each 7T VASO scan is available in Supplementary Table 4. The procedure that we followed, averaging the fMRI response across voxels in a layer ROI that was defined on the upsampled grid, is analogous to taking a weighted average across voxels in the original space (weighted by the proportion of the voxel’s volume that intersects the cortical surface, see Supplementary Fig. 10). The number of voxels in the original resolution in each 7T VASO scan is also available in Supplementary Table 5.

#### Code and sequence availability

Publicly available software packages were used for preprocessing and analysis, including AFNI (https://afni.nimh.nih.gov/) for preprocessing of 3T BOLD data, mrTools (https://github.com/justingardner/mrTools) for preprocessing of 7T BOLD and 7T VASO data, and LayNii toolbox (https://github.com/layerfMRI/LAYNII) for extracting cortical layers. Details of the 7T sequence and scan parameters are available at GitHub (https://github.com/tinaliutong/sequence). Customized code, source data, high-resolution figures, and computational simulation of different experimental designs are available at GitHub (https://github.com/tinaliutong/layerfmri_AMG_V1).

### Data availability

The datasets generated during the current study are available from the corresponding author to the editors and peer reviewers during the review process. Following acceptance, datasets will be freely and publicly available to readers via figshare repository (doi: 10.6084/m9.figshare.14519127).

## Acknowledgements

This work was supported by the Intramural Research Program of the National Institute of Mental Health (ZIAMH002918; NCT00001360). We acknowledge Dr. Larentius Huber for the VASO sequence used here and for his comments on the manuscript. We thank Peter Bandettini, Marlene Behrmann, Kendrick Kay, Zvi Roth, Kunjan Rana, Fernando Ramirez and the CNaP lab at NIMH for helpful comments.

## Author contributions

T.T.L. conceptualized the layer fMRI study, designed the task paradigm, collected the data, analyzed the data, generated figures and wrote the original draft. J.Z.F. collected the data, administered the project, implemented algorithms and generated portions of figures. Y.C. optimized fMRI data acquisition and analysis methodology, wrote portions of the original manuscript and edited the manuscript. S.J. provided the stimuli, contributed to the task paradigm design, and edited the manuscript. G.C. contributed to statistical analyses and visualization, and edited the manuscript. L.G.U. conceptualized the layer fMRI study, supervised study design and interpretation and edited the manuscript (last edit dated: 10/25/2020 with frequent discussions throughout Nov 2020). E.P.M. conceptualized both layer fMRI and BOLD fMRI study, designed the task paradigm, visualized the data, contributed to the analysis methodology, supervised the project and edited the manuscript.

## Supplementary Figures

Supplementary Figure 1 | Average of 3T BOLD (n=11) and 7T BOLD (n=12) participants’ fMRI response to fearful - neutral faces (negative valence effect).

Supplementary Figure 2 | fMRI response to fearful - neutral faces (negative valence effect) on the inflated cortical surface in each 3T BOLD participant who was also scanned in the face localizer experiment (n=11).

Supplementary Figure 3 | fMRI response to fearful - neutral faces (negative valence effect) on the inflated cortical surface in each 7T BOLD participant who was also scanned in the face localizer experiment (n=12).

Supplementary Figure 4 | Assessment of the BML model fit.

Supplementary Figure 5 | Percent change in VASO (in units of ml per 100 ml CBV) to each facial expression (fearful, neutral and happy) as a function of relative cortical depth between WM (left) and CSF (right).

Supplementary Figure 6 | Within-subject test-retest reliability of VASO results across scan sessions.

Supplementary Figure 7 | Task performance across experiments and expressions.

Supplementary Figure 8 | Within-subject test-retest reliability of task performance across scan sessions.

Supplementary Figure 9 | fMRI response to fearful and neutral faces as a function of fixation block length.

Supplementary Figure 10 | A visualization of 21 layers defined in upsampled and original resolution of the VASO anatomy.

## Supplementary Table

Supplementary Table 1 | Numerical values of correlation coefficients for fearful face condition in Figure 2. Number of unique participants scanned at 3T BOLD and 7T BOLD who were also scanned in the face localizer experiment: n=15 (see Table 1).

Supplementary Table 2 | Numerical values of correlation coefficients for happy face condition in Figure 2. Number of unique participants scanned at 3T BOLD and 7T BOLD who were also scanned in the face localizer experiment: n=15 (see Table 1).

Supplementary Table 3 | Numerical values of correlation coefficients for neutral face condition in Figure 2. Number of unique participants scanned at 3T BOLD and 7T BOLD who were also scanned in the face localizer experiment: n=15 (see Table 1).

Supplementary Table 4 | Number of voxels per layer (in the upsampled resolution) in each 7T VASO scan (number of unique participants = 10, total scan session = 15).

Supplementary Table 5 | Number of voxels (in the original resolution) in each 7T VASO scan (number of unique participants = 10, total scan session = 15).

