## Supplementary Materials for "Layer-specific, retinotopically-diffuse modulation in human visual cortex by emotional faces"

**Supplementary Methods**

In all of the fMRI experiments in this study, 18 s blocks of faces with different facial expressions were interspersed with 9 s blocks of fixation. This relatively short fixation block compared to the stimulus blocks did not allow enough time for the hemodynamic response to fully return to baseline. However, because the fMRI response is a shift-invariant linear system, and because the face block order was fully randomized, it was possible to use deconvolution to separately measure the response to each face condition. To confirm that the experimental design was appropriate, we ran a ground-truth simulation in which we generated synthetic fMRI time series using the same experimental design as in the main experiment (with 18 s stimulation blocks and 9 s fixation blocks in between). We first generated a random sequence of trials following the experimental design described in the main experiment (18 s face blocks with conditions for neutral, happy, and fearful conditions) with 9 s fixation blocks interspersed. We then convolved this sequence with a canonical hemodynamic response function (a difference of gammas, following the conventions in SPM). We then added random noise from a Gaussian distribution with mean of zero. The SD of the noise was a parameter that we adjusted. The simulation results in Supplementary Fig. 9a were generated with a signal amplitude of 1, 1.2, and 1.4 for neutral, happy, and fearful conditions. The noise SD was set to 0.4. We found that we could reliably recover the ground-truth fMRI response with the same deconvolution code that we used to analyze the empirical fMRI data (see Supplementary Fig. 9a, left). We repeated the simulation with longer fixation blocks (Supplementary Fig. 9a, middle and right), holding the number of time points in the simulated time series constant. With these longer fixation blocks, we could recover the ground-truth fMRI response, but the variance of the estimate increased, indicating that the shorter 9 s blocks yielded more reliable response estimates. The code implementing this simulation can be found on the GitHub link associated with this manuscript (<https://github.com/tinaliutong/layerfmri_AMG_V1/blob/main/simExptDesign.m>).

We next wanted to confirm these simulations using empirical fMRI data. We collected data from three additional participants across four scanning sessions. Participants were scanned at 7T using BOLD fMRI with the same range of fixation block durations as in the simulation. In addition to the 9 s fixation block design reported in the main text, we included an intermediate-length fixation block design (18 s) and a very long fixation block design (36 s). Each run consisted of three repeats of each facial expression condition (fearful, neutral, happy) in randomized order and ten repeats of the fixation block. In each of the three experimental designs that we tested, we kept the total duration of the face blocks the same across designs. We also kept the total scan time devoted to each experimental design nearly the same, as in the simulation. By testing a range of fixation block durations, we confirmed that the facial valence effects in V1 are robust regardless of the duration of the fixation blocks (Supplementary Fig. 9b).

**Supplementary Figures**

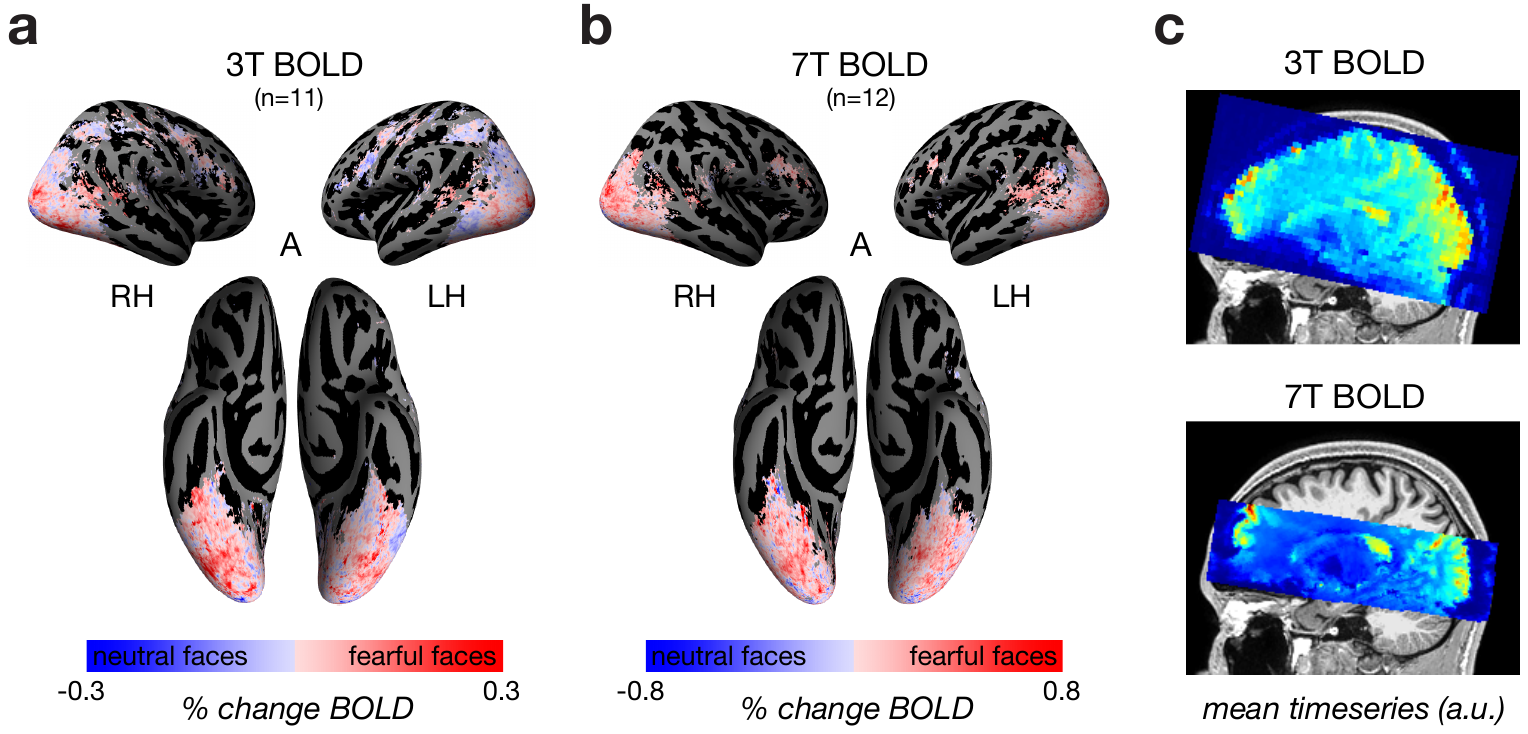

### Supplementary Figure 1 | Average of 3T BOLD (*n*=11) and 7T BOLD (*n*=12) participants’ fMRI response to fearful - neutral faces (negative valence effect). a-b, Lateral (top) and ventral (bottom) views of the fsaverage inflated cortical surface. Superimposed on the cortical surface (color) is the subtraction of response amplitude to fearful faces and neutral faces for each voxel that exhibited a reliable visual response (coefficient of determination R^2^>0.1) in at least one third of the participants. Stronger response to fearful faces (light and dark red); stronger response to neutral faces (light and dark blue). c, Field of view in 3T BOLD and 7T BOLD experiments, indicated by mean timeseries of a single run from the same participant (scaled min to max). 3T BOLD protocol covered entire cortex; 7T BOLD experiment focused on visual cortex and the amygdala.

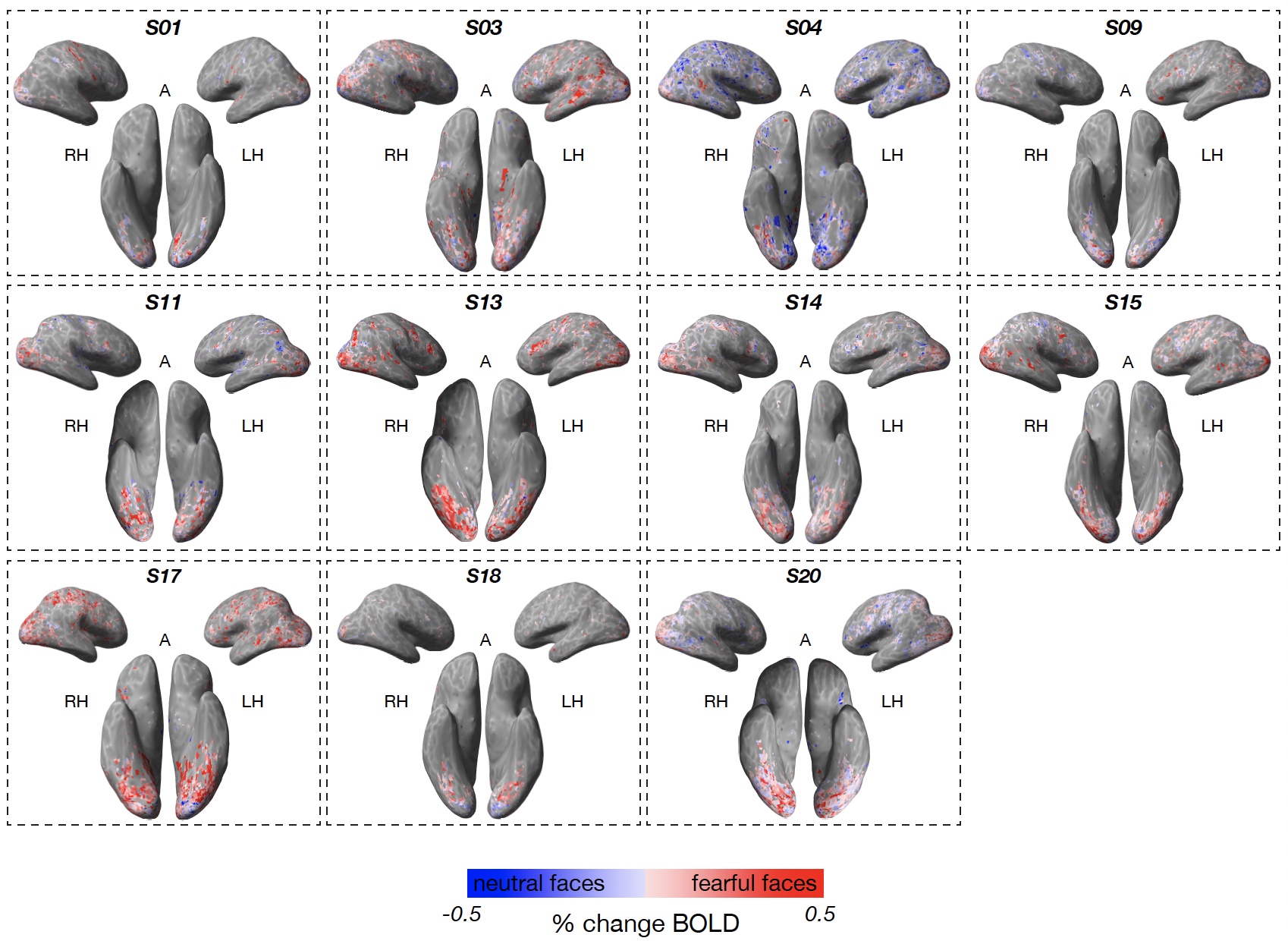

**Supplementary Figure 2 | fMRI response to fearful - neutral faces (negative** **valence effect) on the inflated cortical surface in each 3T BOLD participant who was also scanned in the face localizer experiment (*n*=11).** Lateral (top) and ventral (bottom) views of each participant’s inflated cortical surface in native space. Superimposed on the cortical surface (color) is the subtraction of response amplitude to fearful faces and neutral faces for each voxel that exhibited a reliable visual response (R^2^>0.1). Stronger response to fearful faces (light and dark red); stronger response to neutral faces (light and dark blue).

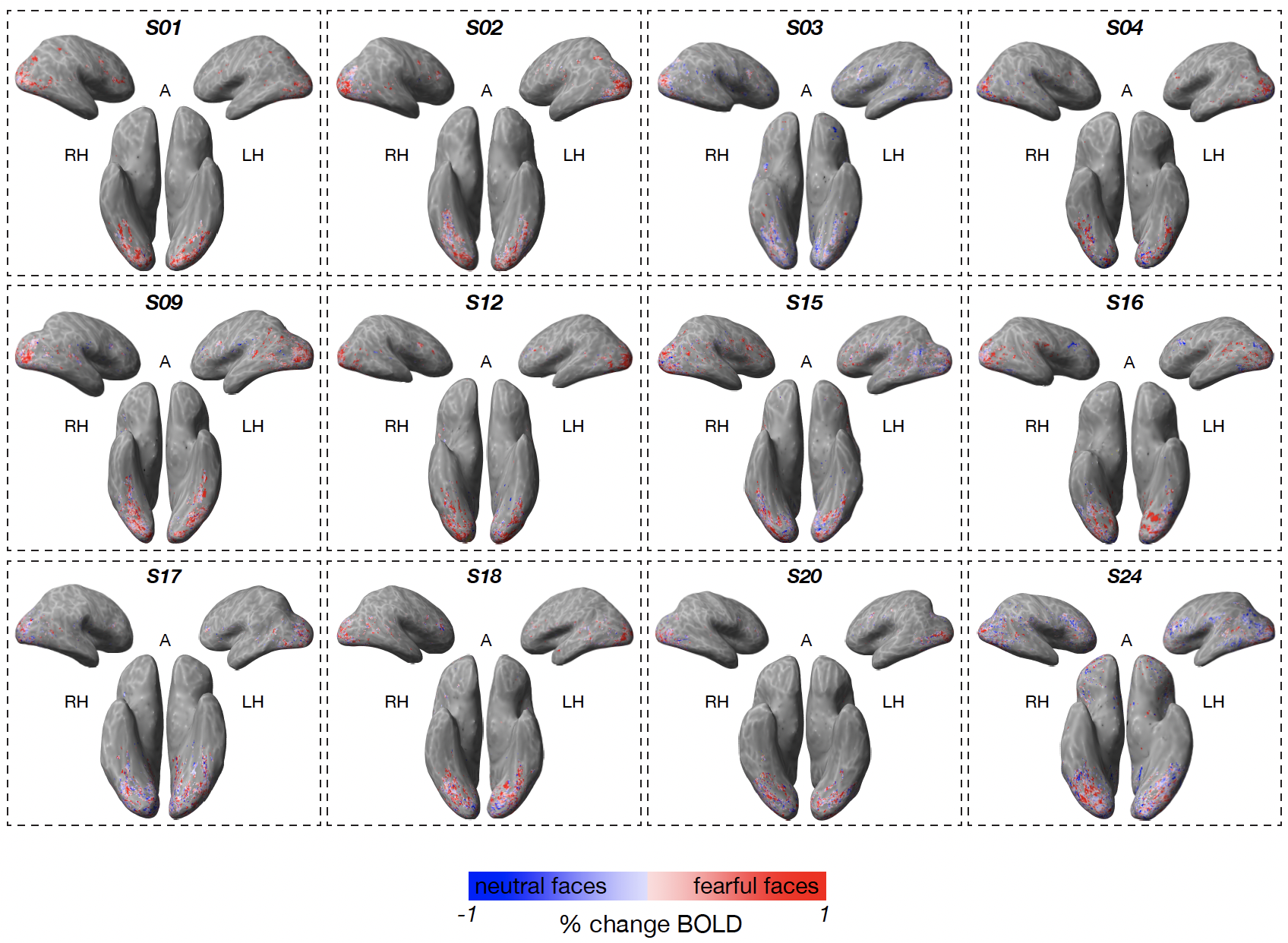

**Supplementary Figure 3 | fMRI response to fearful - neutral faces (negative** **valence effect) on the inflated cortical surface in each 7T BOLD participant who was also scanned in the face localizer experiment (*n*=12).** Lateral (top) and ventral (bottom) views of each 7T BOLD participant’s inflated cortical surface in native space. Superimposed on the cortical surface (color) is the subtraction of response amplitude to fearful faces and neutral faces for each voxel that exhibited a reliable visual response (coefficient of determination R^2^>0.1). Stronger response to fearful faces (light and dark red); stronger response to neutral faces (light and dark blue).

**Supplementary Figure 4 | Assessment of the BML model fit.** The accuracy and adequacy of the adopted BML model assessed through posterior predictive checks overlaid with the raw data. The solid black curve is the kernel density of the raw data with linear interpolation while the fat curve in light blue is composed of 200 sub-curves each of which corresponds to one draw from the posterior distribution based on the BML model. The differences between these two curves indicate how well the BML model fits the raw data.

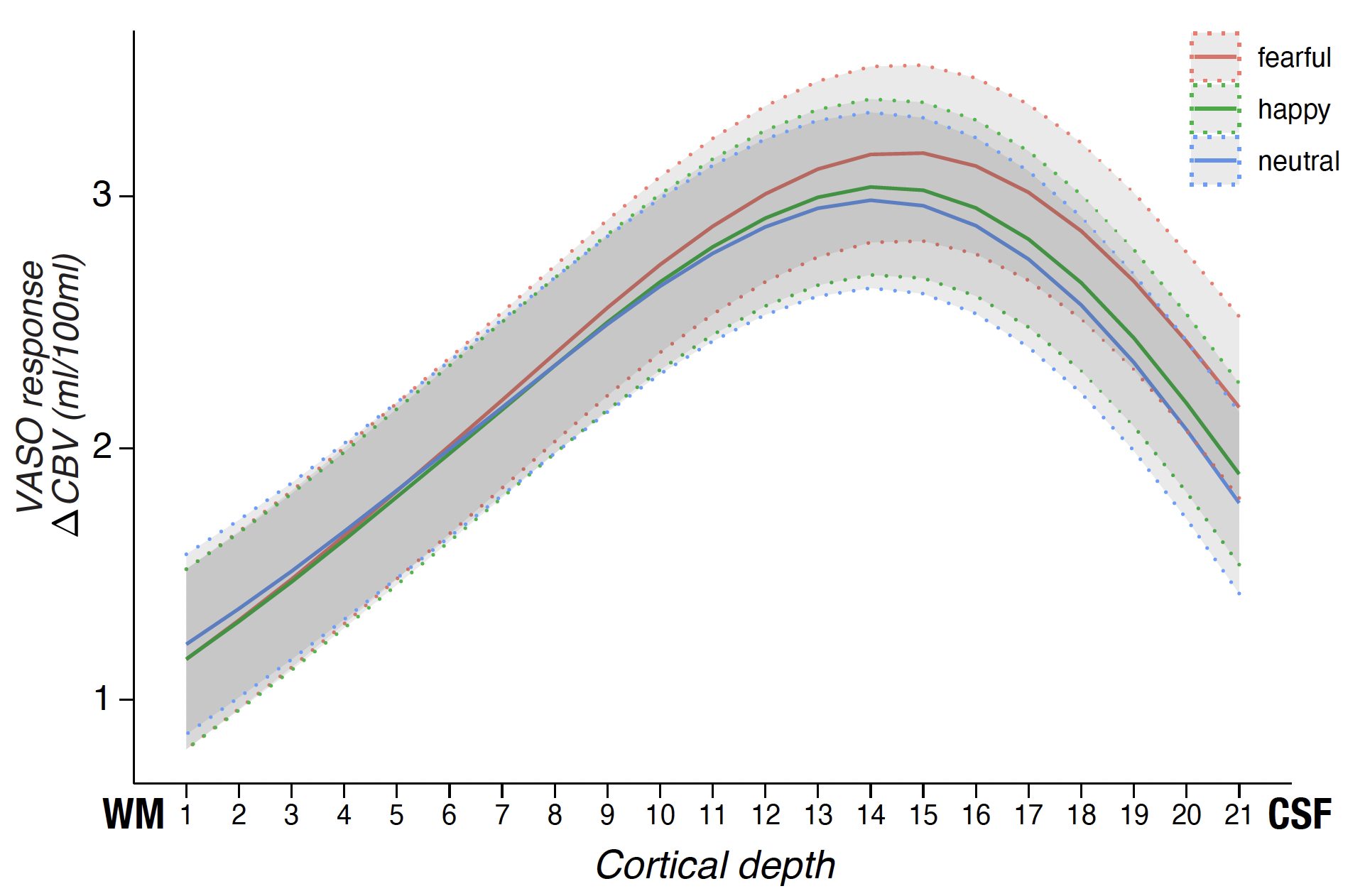

**Supplementary Figure 5 | Percent change in VASO (in units of ml per 100 ml CBV) to each facial expression (fearful, neutral, and happy) as a function of relative cortical depth between WM (left) and CSF (right).** Red line, fearful; green line, happy; blue line, neutral. The shaded bands bounded with dotted lines indicate the uncertainty range of one standard error for the three fitted profiles. Number of unique participants scanned at 7T VASO: *n*=10 (15 scan sessions, see Table 1).

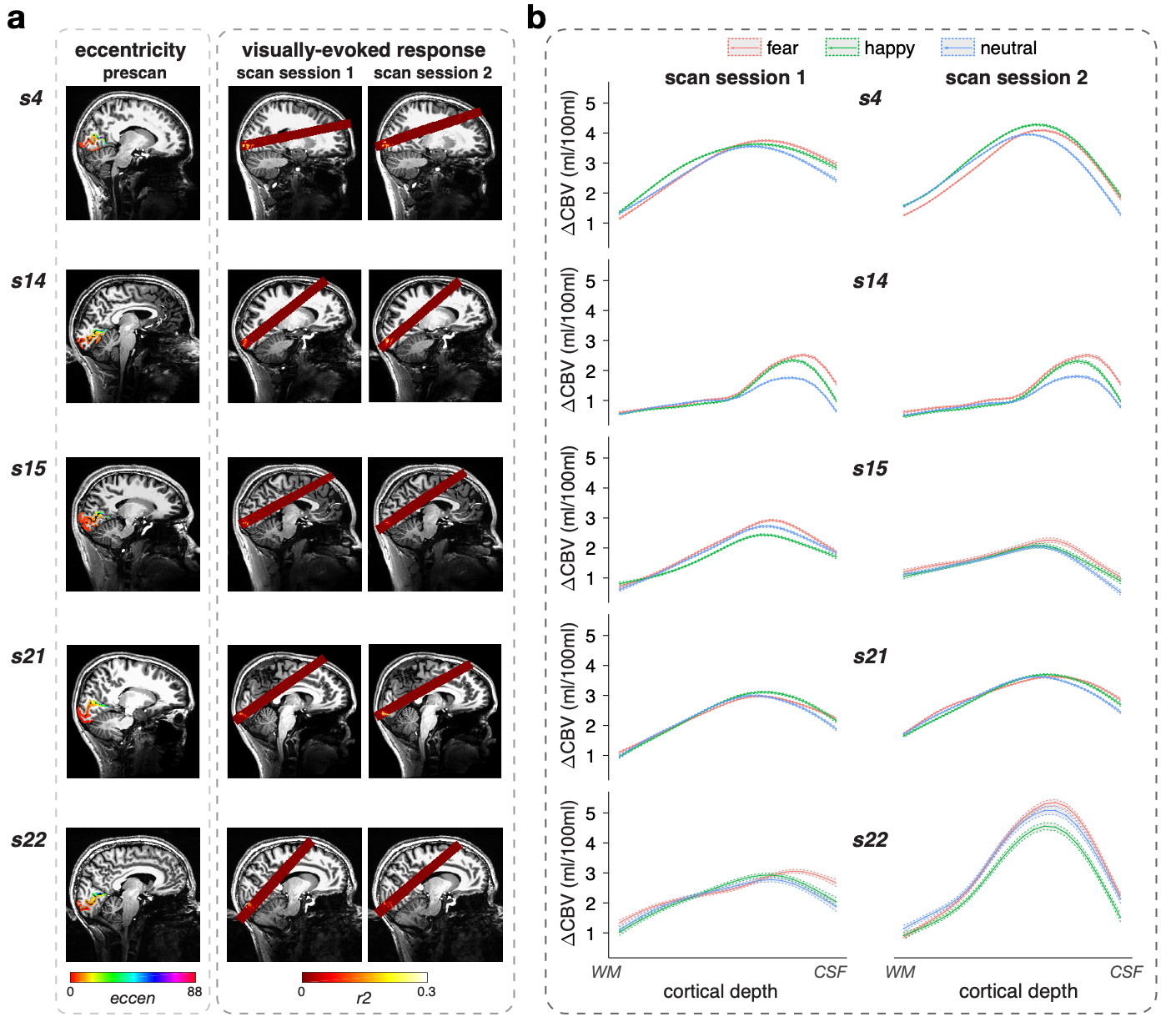

**Supplementary Figure 6 | Within-subject test-retest reliability of VASO results across scan sessions. a**, A visual eccentricity map (left) was visualized based on each participant’s structural MRI scan acquired in a separate session prior to the main experimental scan session, which was used to guide slice prescription, aiming to maximally cover V1 in each participant and in each session (right). **b**, Percent change in VASO (in units of ml per 100 ml CBV) to each facial expression (fearful, neutral and happy) as a function of relative cortical depth between WM (left) and CSF (right). Red line, fearful; green line, happy; blue line, neutral. Each shaded band bounded with dotted lines indicates the uncertainty range of one standard error for each cross-layer profile. Left, scan session 1; right, scan session 2. Participant numbers here correspond to those in Table 1.

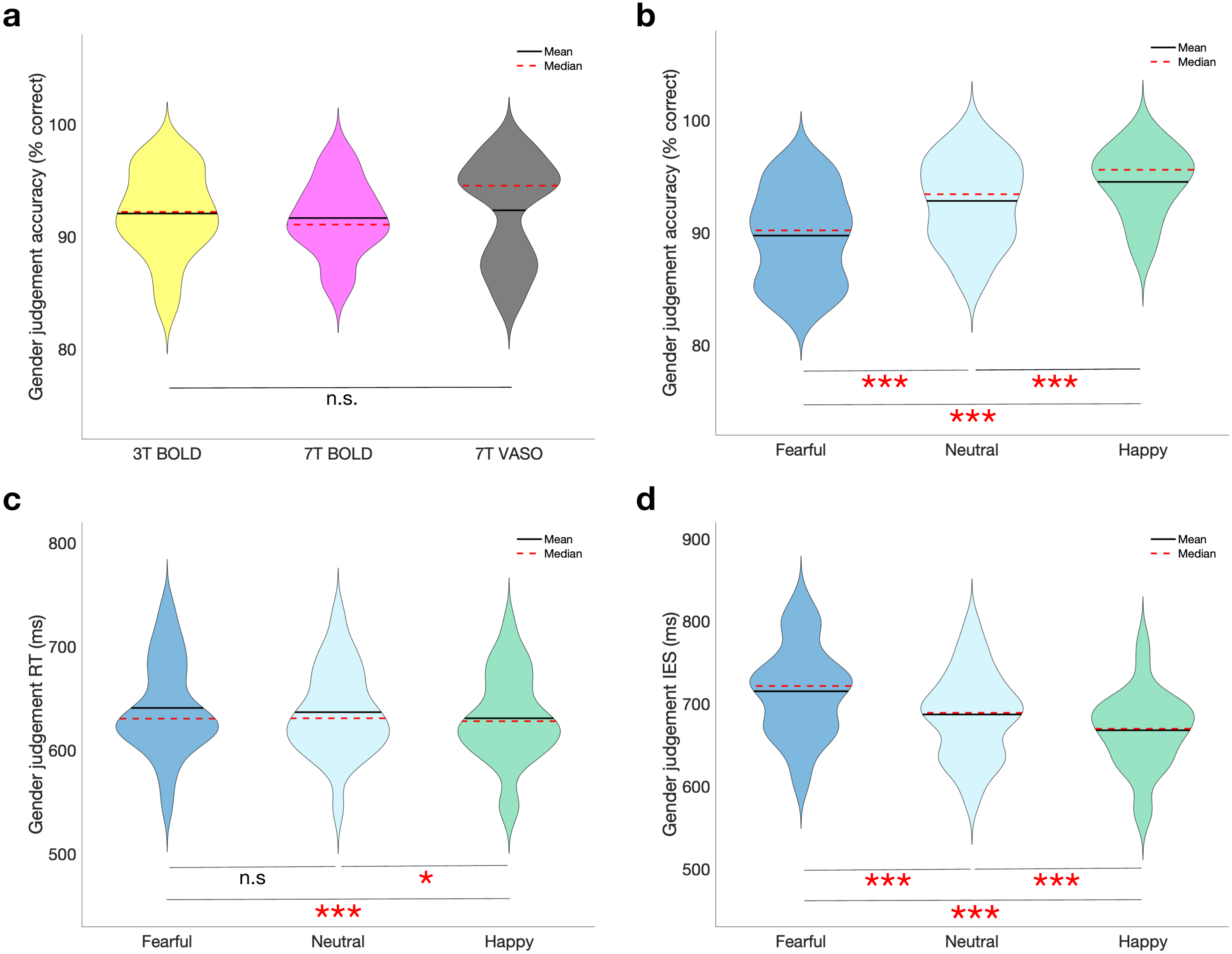

**Supplementary Figure 7 | Task performance across experiments and expressions.**

**a**, Accuracy in the gender judgment task (collapsed across expression) did not differ across the three experiments. Each violin plot shows the full distribution of accuracy with black solid line indicating the mean and red dashed line indicating the median. 7T VASO: n = 10 unique participants (15 scan sessions), 7T BOLD: n = 14 participants (14 scan sessions), 3T BOLD: n = 14 participants (14 scan sessions). One-way ANOVA: *F*(2,35) = .094, *P* = 0.910, n.s., not significant. **b**, Accuracy in the gender judgment task revealed a significant difference across expressions. Each violin plot shows the full distribution of accuracy with black solid line indicating the mean and red dashed line indicating the median. One-way repeated measures ANOVA: *F*(2,48) = 51.520, *P* < 0.001, partial *η*^2^ = 0.682. **c**, Reaction time (RT) in the gender judgment task revealed a significant difference across expressions. Each violin plot shows the full distribution of RTs (correct trials) with black solid line indicating the mean and red dashed line indicating the median. One-way repeated measures ANOVA: *F*(2,48) = 8.906, *P* < 0.001, partial *η*^2^ = 0.271. **d**, Inverse efficiency score (IES) in the gender judgment task revealed a significant difference across expressions. Each violin plot shows the full distribution of IESs (correct trials) with black solid line indicating the mean and red dashed line indicating the median. One-way repeated measures ANOVA: *F*(2,48) = 68.339, *P* < 0.001, partial *η*^2^ = 0.740. N = 25 participants for each condition (fearful: dark blue, neutral: light blue, happy: green) in **b-d**. *** *P* < 0.001, ** *P* < 0.01, **P* < 0.05, two-tailed paired t-test (in **b-d)**; n.s., not significant.

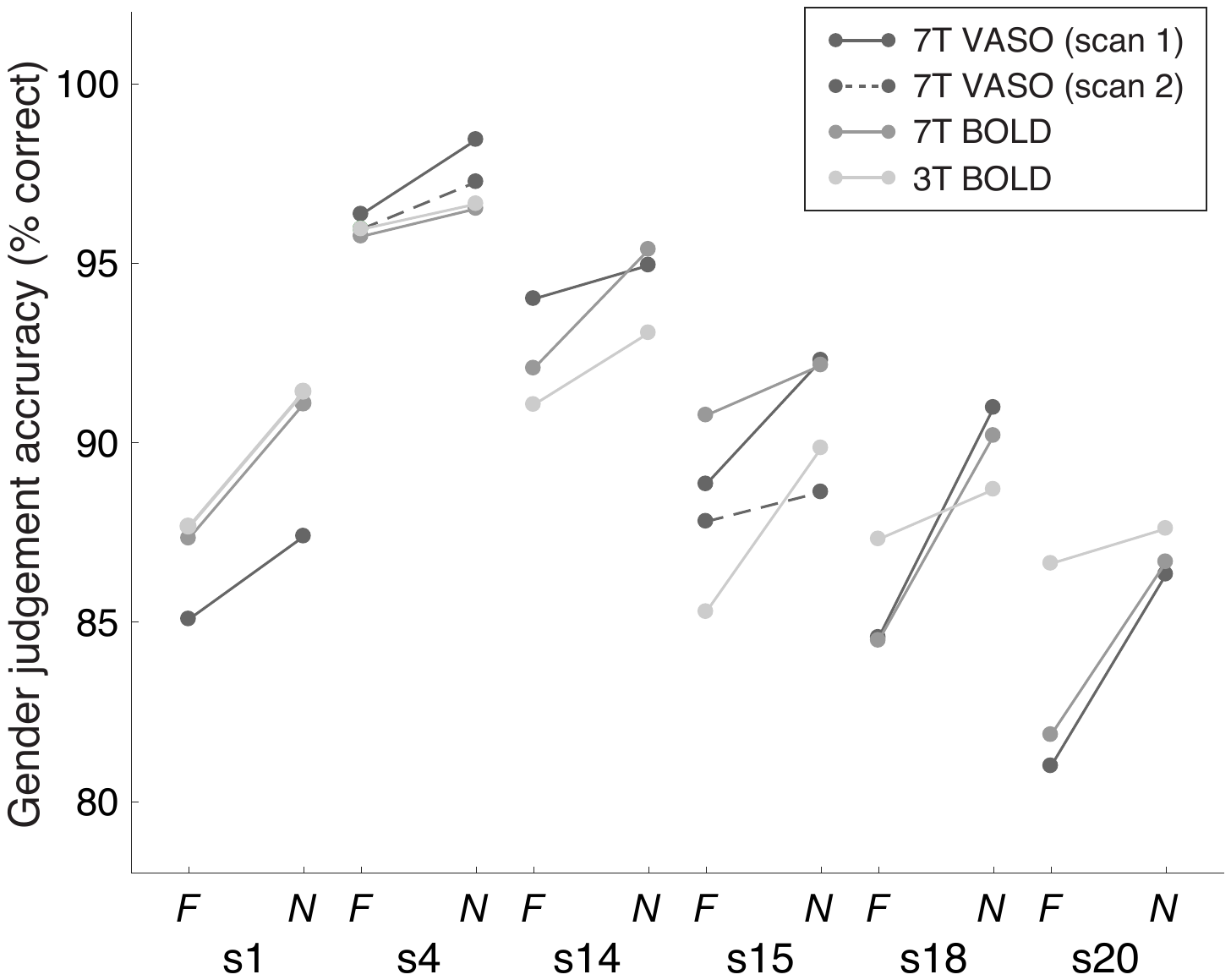

**Supplementary Figure 8 | Within-subject test-retest reliability of task performance across scan sessions.** Performance in the gender judgement task (percent correct) from 6 participants who were scanned in at least three scan sessions. Participant numbers here correspond to those in Table 1. 7T VASO (scan 1): dark gray solid line, 7T VASO (scan 2): dark gray dashed line, 7T BOLD: medium gray, 3T BOLD: light gray. N: neutral, F: fearful.

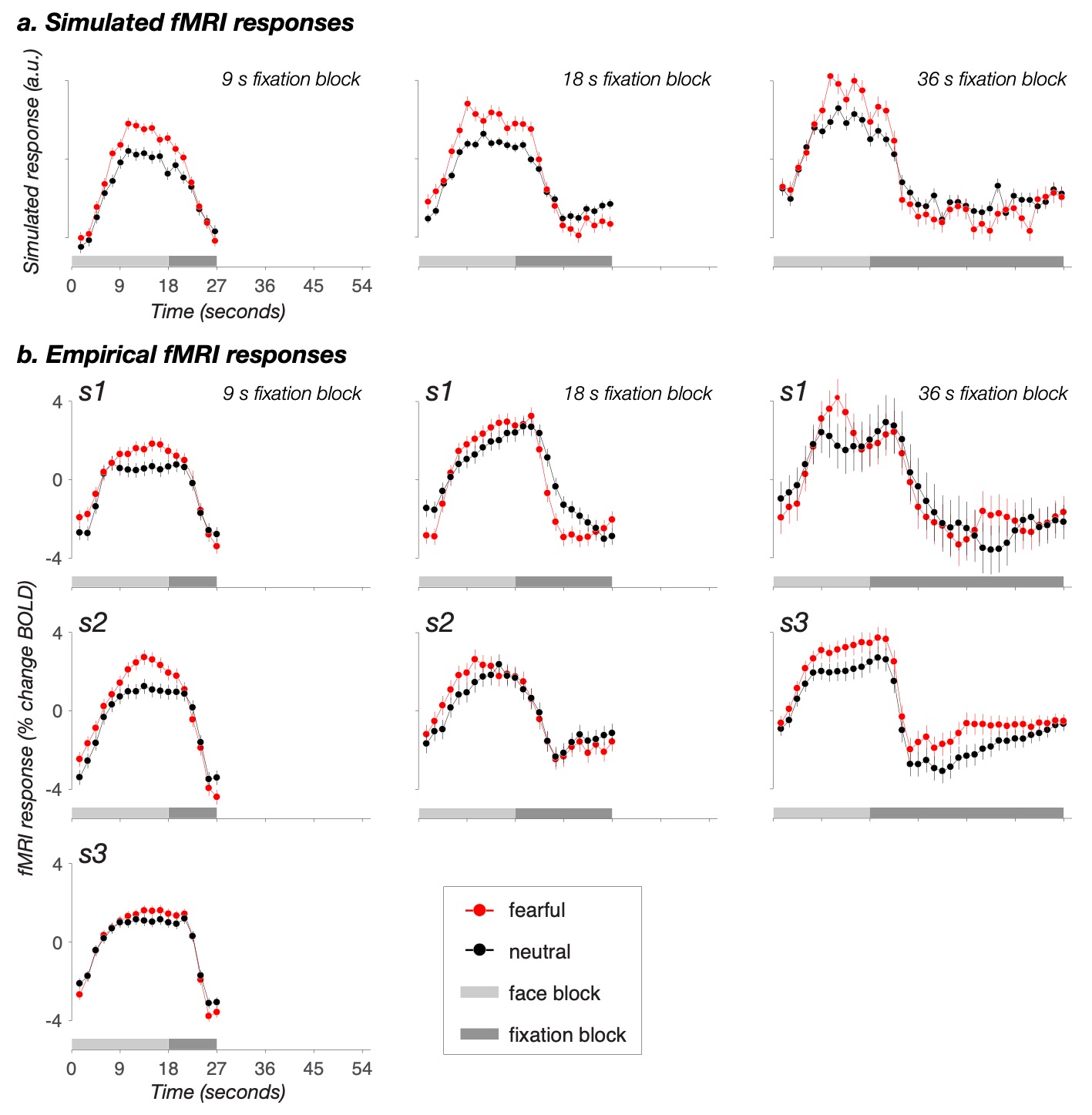

**Supplementary Figure 9 | fMRI response to fearful and neutral faces as a function of fixation block length. a,** Simulated fMRI responses. **b,** Empirical fMRI responses. Left: 9s fixation block design, Middle: 18s fixation block design, Right: 36 s fixation block design. Red line: fearful face condition, Black line: neutral face condition, Light gray box: duration of each face block (18s), Dark gray box: duration of each fixation block (9 s, 18 s, or 36 s). These three participants were scanned as part of the revision process. The data from these participants were not included in the main manuscript and the subject numbers here do not correspond to those in Table 1.

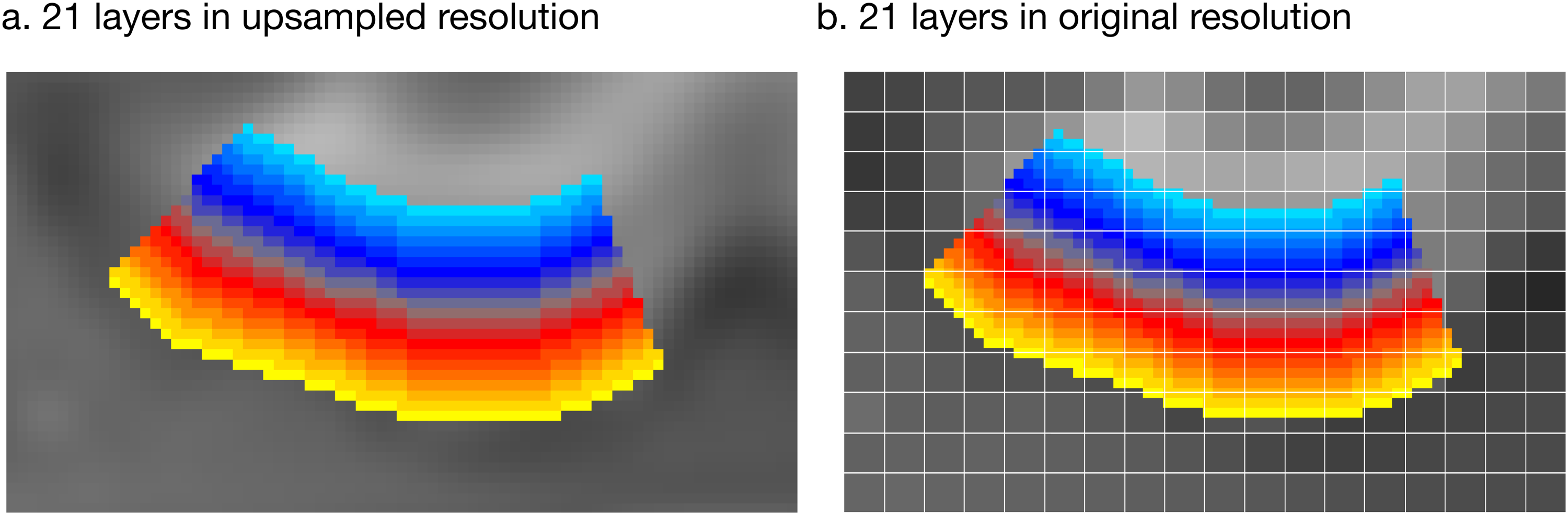

**Supplementary Figure 10 | A visualization of 21 layers defined in upsampled and original resolution of the VASO anatomy. a**, 21 layers in parafoveal V1 visualized in the upsampled resolution of the VASO anatomy. **b**, the same 21 layers projected back to VASO anatomy in the original resolution.

**Supplementary Tables**

**Supplementary Table 1 | Numerical values of correlation coefficients for fearful face condition in Figure 2. Number of unique participants scanned at 3T BOLD and 7T BOLD who were also scanned in the face localizer experiment: n=15 (see Table 1).**

| 1.000 | 0.238 | 0.297 | 0.265 | 0.279 | 0.298 | 0.360 | 0.309 | 0.304 | 0.313 | 0.374 | 0.292 | 0.324 | 0.281 |
| --- | --- | --- | --- | --- | --- | --- | --- | --- | --- | --- | --- | --- | --- |
| 0.238 | 1.000 | 0.758 | 0.882 | 0.848 | 0.707 | 0.554 | 0.574 | 0.363 | 0.637 | 0.493 | 0.661 | 0.495 | 0.442 |
| 0.297 | 0.758 | 1.000 | 0.747 | 0.694 | 0.592 | 0.472 | 0.598 | 0.465 | 0.555 | 0.532 | 0.642 | 0.509 | 0.425 |
| 0.265 | 0.882 | 0.747 | 1.000 | 0.929 | 0.761 | 0.583 | 0.648 | 0.420 | 0.710 | 0.603 | 0.745 | 0.551 | 0.455 |
| 0.280 | 0.848 | 0.694 | 0.929 | 1.000 | 0.827 | 0.669 | 0.708 | 0.452 | 0.796 | 0.644 | 0.800 | 0.603 | 0.516 |
| 0.298 | 0.707 | 0.592 | 0.761 | 0.827 | 1.000 | 0.726 | 0.771 | 0.487 | 0.772 | 0.607 | 0.697 | 0.640 | 0.542 |
| 0.360 | 0.554 | 0.472 | 0.583 | 0.669 | 0.726 | 1.000 | 0.625 | 0.418 | 0.723 | 0.626 | 0.572 | 0.628 | 0.568 |
| 0.309 | 0.574 | 0.598 | 0.648 | 0.708 | 0.771 | 0.625 | 1.000 | 0.665 | 0.728 | 0.655 | 0.749 | 0.694 | 0.552 |
| 0.304 | 0.363 | 0.465 | 0.420 | 0.452 | 0.487 | 0.418 | 0.665 | 1.000 | 0.514 | 0.509 | 0.537 | 0.547 | 0.432 |
| 0.313 | 0.637 | 0.555 | 0.710 | 0.796 | 0.772 | 0.723 | 0.728 | 0.514 | 1.000 | 0.747 | 0.770 | 0.738 | 0.602 |
| 0.374 | 0.493 | 0.532 | 0.603 | 0.644 | 0.607 | 0.626 | 0.655 | 0.509 | 0.747 | 1.000 | 0.702 | 0.688 | 0.574 |
| 0.292 | 0.661 | 0.642 | 0.745 | 0.800 | 0.697 | 0.572 | 0.749 | 0.537 | 0.770 | 0.702 | 1.000 | 0.726 | 0.550 |
| 0.324 | 0.495 | 0.509 | 0.551 | 0.603 | 0.640 | 0.628 | 0.694 | 0.547 | 0.738 | 0.688 | 0.726 | 1.000 | 0.739 |
| 0.281 | 0.442 | 0.425 | 0.455 | 0.516 | 0.542 | 0.568 | 0.552 | 0.432 | 0.602 | 0.574 | 0.550 | 0.739 | 1.000 |

**Supplementary Table 2 | Numerical values of correlation coefficients for happy face condition in Figure 2. Number of unique participants scanned at 3T BOLD and 7T BOLD who were also scanned in the face localizer experiment: n=15 (see Table 1).**

| 1.000 | 0.233 | 0.291 | 0.248 | 0.260 | 0.281 | 0.334 | 0.300 | 0.276 | 0.291 | 0.328 | 0.267 | 0.319 | 0.276 |
| --- | --- | --- | --- | --- | --- | --- | --- | --- | --- | --- | --- | --- | --- |
| 0.233 | 1.000 | 0.768 | 0.888 | 0.854 | 0.700 | 0.549 | 0.557 | 0.343 | 0.633 | 0.496 | 0.651 | 0.495 | 0.450 |
| 0.291 | 0.768 | 1.000 | 0.747 | 0.692 | 0.575 | 0.461 | 0.586 | 0.452 | 0.542 | 0.523 | 0.629 | 0.530 | 0.437 |
| 0.248 | 0.888 | 0.747 | 1.000 | 0.933 | 0.740 | 0.566 | 0.629 | 0.405 | 0.697 | 0.594 | 0.735 | 0.555 | 0.461 |
| 0.260 | 0.854 | 0.692 | 0.933 | 1.000 | 0.820 | 0.659 | 0.697 | 0.447 | 0.795 | 0.634 | 0.794 | 0.607 | 0.510 |
| 0.281 | 0.700 | 0.575 | 0.740 | 0.820 | 1.000 | 0.724 | 0.754 | 0.472 | 0.762 | 0.593 | 0.680 | 0.639 | 0.531 |
| 0.334 | 0.549 | 0.461 | 0.566 | 0.659 | 0.724 | 1.000 | 0.606 | 0.411 | 0.725 | 0.614 | 0.565 | 0.626 | 0.549 |
| 0.300 | 0.557 | 0.586 | 0.629 | 0.697 | 0.754 | 0.606 | 1.000 | 0.679 | 0.726 | 0.646 | 0.748 | 0.705 | 0.549 |
| 0.276 | 0.343 | 0.452 | 0.405 | 0.447 | 0.472 | 0.411 | 0.679 | 1.000 | 0.510 | 0.500 | 0.549 | 0.565 | 0.413 |
| 0.291 | 0.633 | 0.542 | 0.697 | 0.795 | 0.762 | 0.725 | 0.726 | 0.510 | 1.000 | 0.740 | 0.768 | 0.748 | 0.586 |
| 0.328 | 0.496 | 0.523 | 0.594 | 0.634 | 0.593 | 0.614 | 0.646 | 0.500 | 0.740 | 1.000 | 0.690 | 0.693 | 0.564 |
| 0.267 | 0.651 | 0.629 | 0.735 | 0.794 | 0.680 | 0.565 | 0.748 | 0.549 | 0.768 | 0.690 | 1.000 | 0.740 | 0.542 |
| 0.319 | 0.495 | 0.530 | 0.555 | 0.607 | 0.639 | 0.626 | 0.705 | 0.565 | 0.748 | 0.693 | 0.740 | 1.000 | 0.730 |
| 0.276 | 0.450 | 0.437 | 0.461 | 0.510 | 0.531 | 0.549 | 0.549 | 0.413 | 0.586 | 0.564 | 0.542 | 0.730 | 1.000 |

**Supplementary Table 3 | Numerical values of correlation coefficients for neutral face condition in Figure 2. Number of unique participants scanned at 3T BOLD and 7T BOLD who were also scanned in the face localizer experiment: n=15 (see Table 1).**

| 1.000 | 0.197 | 0.258 | 0.221 | 0.231 | 0.239 | 0.309 | 0.258 | 0.246 | 0.271 | 0.301 | 0.236 | 0.297 | 0.252 |
| --- | --- | --- | --- | --- | --- | --- | --- | --- | --- | --- | --- | --- | --- |
| 0.197 | 1.000 | 0.754 | 0.879 | 0.845 | 0.696 | 0.524 | 0.553 | 0.338 | 0.623 | 0.470 | 0.644 | 0.461 | 0.442 |
| 0.258 | 0.754 | 1.000 | 0.739 | 0.676 | 0.566 | 0.450 | 0.577 | 0.449 | 0.530 | 0.514 | 0.624 | 0.512 | 0.439 |
| 0.221 | 0.879 | 0.739 | 1.000 | 0.921 | 0.739 | 0.546 | 0.618 | 0.393 | 0.679 | 0.571 | 0.727 | 0.524 | 0.444 |
| 0.231 | 0.845 | 0.676 | 0.921 | 1.000 | 0.821 | 0.635 | 0.693 | 0.429 | 0.777 | 0.609 | 0.781 | 0.581 | 0.490 |
| 0.239 | 0.696 | 0.566 | 0.739 | 0.821 | 1.000 | 0.700 | 0.749 | 0.458 | 0.760 | 0.571 | 0.671 | 0.611 | 0.508 |
| 0.309 | 0.524 | 0.450 | 0.546 | 0.635 | 0.700 | 1.000 | 0.585 | 0.395 | 0.711 | 0.590 | 0.550 | 0.596 | 0.532 |
| 0.258 | 0.553 | 0.577 | 0.618 | 0.693 | 0.749 | 0.585 | 1.000 | 0.673 | 0.726 | 0.627 | 0.749 | 0.696 | 0.549 |
| 0.246 | 0.338 | 0.449 | 0.393 | 0.429 | 0.458 | 0.395 | 0.673 | 1.000 | 0.517 | 0.507 | 0.549 | 0.567 | 0.421 |
| 0.271 | 0.623 | 0.530 | 0.679 | 0.777 | 0.760 | 0.711 | 0.726 | 0.517 | 1.000 | 0.714 | 0.746 | 0.724 | 0.587 |
| 0.301 | 0.470 | 0.514 | 0.571 | 0.609 | 0.571 | 0.590 | 0.627 | 0.507 | 0.714 | 1.000 | 0.670 | 0.675 | 0.553 |
| 0.236 | 0.644 | 0.624 | 0.727 | 0.781 | 0.671 | 0.550 | 0.749 | 0.549 | 0.746 | 0.670 | 1.000 | 0.722 | 0.537 |
| 0.297 | 0.461 | 0.512 | 0.524 | 0.581 | 0.611 | 0.596 | 0.696 | 0.567 | 0.724 | 0.675 | 0.722 | 1.000 | 0.726 |
| 0.252 | 0.442 | 0.439 | 0.444 | 0.490 | 0.508 | 0.532 | 0.549 | 0.421 | 0.587 | 0.553 | 0.537 | 0.726 | 1.000 |

**Supplementary Table 4 | Number of voxels per layer (in the upsampled resolution) in each 7T VASO scan (number of unique participants = 10, total scan session = 15).**

| Participant | **Scan session** | **Layer** | | | | | | | | | | | | | | | | | | | | |
| --- | --- | --- | --- | --- | --- | --- | --- | --- | --- | --- | --- | --- | --- | --- | --- | --- | --- | --- | --- | --- | --- | --- |
|  |  | **1** | **2** | **3** | **4** | **5** | **6** | **7** | **8** | **9** | **10** | **11** | **12** | **13** | **14** | **15** | **16** | **17** | **18** | **19** | **20** | **21** |
| 1 | **1** | 251 | 245 | 293 | 229 | 265 | 215 | 217 | 226 | 220 | 223 | 217 | 240 | 215 | 217 | 223 | 217 | 243 | 223 | 296 | 290 | 271 |
| 4 | **1** | 258 | 232 | 218 | 213 | 220 | 221 | 220 | 218 | 227 | 223 | 235 | 236 | 261 | 253 | 277 | 266 | 275 | 283 | 286 | 267 | 301 |
|  | **2** | 271 | 242 | 229 | 218 | 229 | 235 | 232 | 219 | 233 | 236 | 251 | 237 | 263 | 251 | 261 | 259 | 262 | 269 | 279 | 271 | 300 |
| 10 | **1** | 287 | 247 | 257 | 232 | 216 | 239 | 214 | 227 | 218 | 233 | 220 | 245 | 236 | 245 | 235 | 274 | 245 | 271 | 288 | 238 | 261 |
| 14 | **1** | 412 | 414 | 417 | 421 | 386 | 417 | 373 | 381 | 397 | 327 | 371 | 349 | 373 | 329 | 365 | 365 | 393 | 358 | 405 | 363 | 398 |
|  | **2** | 384 | 386 | 391 | 408 | 386 | 413 | 375 | 354 | 368 | 312 | 355 | 334 | 352 | 312 | 334 | 322 | 338 | 318 | 346 | 309 | 338 |
| 15 | **1** | 259 | 229 | 237 | 229 | 253 | 241 | 269 | 263 | 271 | 297 | 310 | 329 | 332 | 322 | 317 | 325 | 322 | 319 | 307 | 309 | 298 |
|  | **2** | 277 | 238 | 242 | 257 | 244 | 238 | 249 | 238 | 248 | 267 | 270 | 288 | 289 | 287 | 277 | 284 | 289 | 286 | 271 | 290 | 305 |
| 20 | **1** | 239 | 228 | 234 | 230 | 226 | 245 | 223 | 232 | 226 | 241 | 261 | 230 | 255 | 269 | 265 | 299 | 293 | 289 | 297 | 301 | 297 |
| 21 | **1** | 252 | 216 | 257 | 223 | 214 | 221 | 225 | 214 | 219 | 227 | 234 | 214 | 250 | 245 | 234 | 245 | 248 | 266 | 254 | 243 | 261 |
| 22 | **1** | 275 | 256 | 266 | 249 | 224 | 212 | 227 | 216 | 218 | 230 | 245 | 223 | 252 | 248 | 236 | 248 | 250 | 272 | 257 | 248 | 288 |
|  | **2** | 285 | 283 | 265 | 277 | 250 | 237 | 254 | 230 | 228 | 231 | 232 | 219 | 241 | 238 | 237 | 241 | 248 | 257 | 248 | 250 | 281 |
| 23 | **1** | 264 | 245 | 235 | 254 | 257 | 266 | 236 | 236 | 252 | 234 | 275 | 270 | 252 | 261 | 273 | 270 | 305 | 298 | 293 | 287 | 292 |
|  | **2** | 268 | 233 | 249 | 238 | 242 | 249 | 253 | 242 | 259 | 261 | 278 | 273 | 265 | 303 | 284 | 284 | 299 | 336 | 313 | 302 | 311 |
| 24 | **1** | 285 | 271 | 292 | 311 | 307 | 333 | 335 | 321 | 307 | 319 | 344 | 349 | 327 | 324 | 299 | 338 | 326 | 306 | 304 | 324 | 323 |

**Supplementary Table 5 | Number of voxels (in the original resolution) in each 7T VASO scan (number of unique participants = 10, total scan session = 15).**

| **Participant** | **Scan session** | **Number of voxels in the original resolution** |
| --- | --- | --- |
| **1** | 1 | 317 |
| **4** | 1 | 323 |
|  | 2 | 330 |
| **10** | 1 | 322 |
| **14** | 1 | 499 |
|  | 2 | 464 |
| **15** | 1 | 375 |
|  | 2 | 354 |
| **20** | 1 | 334 |
| **21** | 1 | 312 |
| **22** | 1 | 323 |
| **23** | 2 | 328 |
|  | 1 | 345 |
| **24** | 2 | 361 |
|  | 1 | 412 |
